# Limited effect of short- to mid-term storage conditions on an Australian farmland soil RNA virome

**DOI:** 10.1101/2025.08.27.672739

**Authors:** Sabrina Sadiq, PeiPei Xue, Yijia Tang, Mingming Du, Kate Van Brussel, Alex B. McBratney, Edward C. Holmes, Budiman Minasny

## Abstract

Soils represent one of the largest and most diverse reservoirs of microbial life on Earth, yet their associated RNA viruses remain underexplored compared to animal and aquatic systems. Viral discovery in soils has been further limited by technical hurdles, particularly obtaining sufficient yields of high-quality RNA for sequencing. To address this, we evaluated a range of storage and preservation strategies, including the use of commercial preservative solutions and ultra-cold snap-freezing, followed by standardised RNA extraction, sequencing, and virus discovery pipelines. This work aimed to establish minimum sample storage requirements that maintain RNA integrity, generate sufficient RNA sequencing data, and subsequently enable reliable soil virome characterisation. While no preservative solution proved effective, “neat” soil samples were stable at 2–8°C and −30°C for at least two weeks, and at −80°C for at least three months, with no measurable reduction in RNA quality, sequencing data, or viral abundance and diversity. From 32 resulting sequencing libraries, we identified 1,475 putative novel RNA viruses, with the majority belonging to the microbe-associated phylum *Lenarviricota*. Several novel viruses formed divergent clusters with other environmentally derived sequences distantly related to traditionally animal-associated families such as the *Astroviridae* and *Picornaviridae*. Furthermore, unique clusters within the *Picobirnaviridae*, *Alsuvurucetes*, *Ghabrivirales*, and *Amabiliviricetes* comprised exclusively Australian viruses, suggesting instances of region-specific evolution. Together, these findings highlight soils as rich reservoirs of RNA viral diversity and provide practical minimum standards for storage, expanding opportunities to investigate the ecological and evolutionary roles of RNA viruses in terrestrial systems.

**Importance:** RNA viruses are the most abundant and diverse biological entities on Earth and are likely present in all other organisms and ecosystems, including soil-dwelling invertebrates, microbes, and plants. Despite this, their diversity and role in soil systems remains largely unknown. Methodological challenges in preserving and extracting sufficient quantities of RNA from soils have hindered the study of these communities. Here, we identified 1,475 previously undescribed RNA viruses in Australian soils while systematically testing different preservation strategies. The significance of our research lies in the demonstration that snap-freezing soil is a viable and robust storage strategy for at least three months, while also highlighting the extraordinary scale of viral diversity present in terrestrial environments. This work establishes a foundation for reliable exploration of terrestrial RNA viruses, improving the accessibility of more remote environmental viromes and enabling future efforts to integrate them into broader models of microbial ecology and ecosystem function.

## Introduction

Terrestrial environments exhibit rich biodiversity, comprising intricate networks of plants, invertebrates, fungi, bacteria, and protists. These organisms interact in complex ways to regulate decomposition, nutrient cycling, and plant health (1), and microbial community compositions are tightly linked to the physicochemical properties of the surrounding soil (2–4). RNA viruses are widespread among these soil-dwelling hosts, and also infect animals that interact with the soil through diet, waste, decay, and organic debris (5). While viruses are well-studied in aquatic environments due to their roles in nutrient release, microbial population control, and contributions to host survivability and diversity (6–8), soil viromes are comparatively underexplored (9, 10).

Metatranscriptomic sequencing-based approaches have allowed the field of terrestrial RNA virus ecology to progress rapidly by providing a snapshot of all replicating organisms – including RNA viruses – in a sample. These studies have revealed that virome composition depends on such factors as the local biome, geographical location, pH, organic carbon content, and nutrient availability in a soil sample as well as land use, soil type, and depth (11–15). This, in turn, suggests that RNA viruses may play their own roles in maintaining soil health and regulating biogeochemical cycles, likely through interactions with their soil-dwelling hosts. While this has been proposed for DNA viruses (16–20), to date there is limited evidence of RNA viruses directly participating in these processes (11, 21).

Previous studies have revealed huge numbers of viruses in soil, with over 16,000 *Riboviria* sequences from environmental metagenomes available in the non-redundant (nr) protein database on the National Center for Biotechnology Information (NCBI)/GenBank. Microbe-associated RNA viruses, especially from the *Leviviricetes*, *Narnaviridae*, *Mitoviridae*, and *Botourmiaviridae*, are commonly found in soil, as are virus groups traditionally thought to be animal-associated, including the *Astroviridae*, *Picornavirales*, and *Nodamuvirales*, although these usually fall in lineages distant to their animal-infecting relatives (11, 12, 15). In addition, analyses of publicly available environmental data sets, especially those on the Sequence Read Archive (SRA), have provided insights into global patterns of viral diversity (23–25), helped resolve complex evolutionary relationships (22, 26, 27), and revealed virus-host dynamics (28–30). Metatranscriptomic sequencing also provides a viable method for monitoring farmland for RNA viruses that pose a pathogenic risk to crops and which can have devastating economic consequences for the agricultural industry (31–33). With intensifying land use and climate variability, understanding viral reservoirs in soil is critical for anticipating emerging plant disease risks.

Despite the utility of metatranscriptomic sequencing, its utility may be hindered by difficulties in extracting sufficient yields of high-quality RNA from soil samples. Soil RNA is prone to rapid degradation by microbial RNase activity, as well as inherent soil properties such as temperature, pH, metal ions, and humidity-driven hydrolysis or oxidation (34, 35). In addition, virus sequences can easily be overwhelmed by host transcripts, limiting detection and characterisation opportunities (36, 37). The extraction and careful storage of soil samples prior to returning to laboratory facilities, particularly in remote areas where cold storage isn’t feasible for extended time periods (15), is therefore of critical importance. These limitations may skew global virome data toward easily accessed, well-resourced regions, leaving the majority of terrestrial ecosystems untapped. Improving RNA preservation and extraction protocols will enable more inclusive sampling across diverse biomes, expanding the known RNA virosphere and helping to reveal the precise relationships between virome composition and soil properties. Access to remote environmental viromes will also improve global virus surveillance and hence monitoring agricultural risks (5).

Here, we compared the success of RNA extraction, as well as the subsequent levels of viral abundance and diversity, in sequencing results from soil samples collected from two sites in Sydney (New South Wales, Australia) and stored at a variety of temperatures for up to three months. Three commercially available preservative solutions were tested, as well as “neat” soil with no added preservative. We simulated delays between sample collection and long-term storage in laboratory facilities, as well as delays between freezing and extractions, that are commonly experienced in environmental metatranscriptomic studies, followed by consistent extraction and data processing protocols established to work on a variety of sample types, including soil (11, 12, 15, 38). We hypothesise that RNA yields and quality will degrade over time and that warmer temperatures will provide lower viral abundance and diversity than samples snap frozen at colder temperatures. By simulating realistic delays and storage conditions, we aimed to inform fieldwork planning and improve the reproducibility of soil virome studies, providing validated preservation and storage protocols that enable high-quality RNA extraction from soil.

## Results

### RNA extraction success rates and sequencing data generation

Soil samples stored in RNA PowerProtect or RNA*later* (i.e., all those collected in the first sampling time points of September 2022 and February 2023 from Marrickville, NSW) did not yield a quantifiable amount of RNA, such that they were not prepared for sequencing. Although DNA/RNA Shield seemingly produced appropriate concentrations of RNA, further quality control measures indicated that the RNA was too degraded to successfully generate sequencing libraries. In Camden, NSW (April 2023), preservative-free soils were collected simultaneously with samples preserved in DNA/RNA Shield. The use of DNA/RNA Shield once again resulted in degraded RNA unable to generate sequencing libraries. In contrast, the preservative-free soils did yield sufficient RNA and produced high-quality sequencing data, suggesting snap-freezing soil at −80°C as early as possible after sampling reliably preserves RNA better than preservative solutions when using the extraction, library preparation, and sequencing protocols outlined in the methods. The subsequent shorter trial (September 2023, also Camden) of neat soil stored at 2-8°C, −30°C, and −80°C was similarly successful, indicating a window of at least two weeks where soil can be stored at warmer temperatures and still produce viable RNA for sequencing and downstream analysis.

In total, 32 sequencing libraries were constructed on RNA extracted from the soil samples from Camden (located in south-western Sydney, NSW). The first eight libraries (libraries beginning with “L”) represented the longer trial of soil collected in April 2023 and stored at - 80°C for up to 12 weeks. These libraries represent “data set 1” as shown in Table 1. The remaining 24 libraries were generated from soil collected in September 2023 and stored for up to two weeks at 2-8°C, −30°C, and −80°C (libraries beginning with “C”, “T”, and “E”, respectively). For clarity, these libraries were grouped together as “data set 2” (Table 1). Extracted RNA concentrations, RNA integrity, and sequencing data yields are detailed in Fig. S1. Importantly, no statistically significant relationship was identified between these factors and storage conditions.

**Table 1.** Storage conditions, success rates of RNA extraction and sequencing, and data generation of soil samples collected from Camden (data sets 1 and 2) and Marrickville (data sets 3 and 4) NSW across four time points in 2022 and 2023. Data sets are not listed chronologically as the first two collections (September 2022 and February 2023) did not produce sequence data.

| Data set | Collection month | Storage temperatures | Storage time | Preservative solution | RNA extraction | Sequencing data |
| --- | --- | --- | --- | --- | --- | --- |
| 1 | April 2023 | -80°C | 2, 4, 8, 12 weeks | None | Successful | 8 libraries beginning with “L” |
|  |  |  |  | DNA/RNA Shield | Degraded RNA | No data generated |
| 2 | September 2023 | 2-8°C, -30°C, -80°C | 1, 5, 10, 15 days | None | Successful | 24 libraries beginning with “C”, “T”, or “E” |
| 3 | September 2022 | Ambient, -80°C | 7 days | PowerProtect DNA/RNA | No | Not attempted |
| 4 | February 2023 | Ambient, 2-8°C | 3-6 days | DNA/RNA Shield | Degraded RNA | No data generated |
|  |  |  |  | RNAlater | No | Not attempted |

In total, these 32 libraries generated approximately 2.38 billion sequence reads, 10.4 million of which had similarity to reference sequences within the realm *Riboviria*, kingdom *Orthornavirae* (i.e., RNA viruses). These reads were then assembled into more than 7 million contigs, with initial blast results indicating that 12,568 contigs had sequence similarity to known RNA viruses. Subsequent phylogenetic analysis revealed that a portion of these were mis-aligned to reference sequences or too divergent to confidently be assigned to RNA viral taxa, thus reducing the working data set of RNA virus-like sequences on which statistical analyses were conducted to 7,307 contigs. Viral read abundance in data set 1 ranged from 0.07% in L8B to 0.26% in L2A, stored at −80°C for eight and two weeks, respectively (Fig. 1A). The samples comprising data set 2 all had higher abundance in comparison, ranging from 0.28% in T10B, which was extracted after 10 days of storage at −30°C, up to 1.56% in E15A, which was stored for 15 days at −80°C (Fig. 1A). Interestingly, despite having relatively high viral read abundance, E15A also had the lowest Shannon diversity index of 0.46. The highest Shannon diversity index of 1.74 was seen in C15B, which was extracted after 15 days of storage at 2-8°C (Fig. 1B).

**Figure 1.**
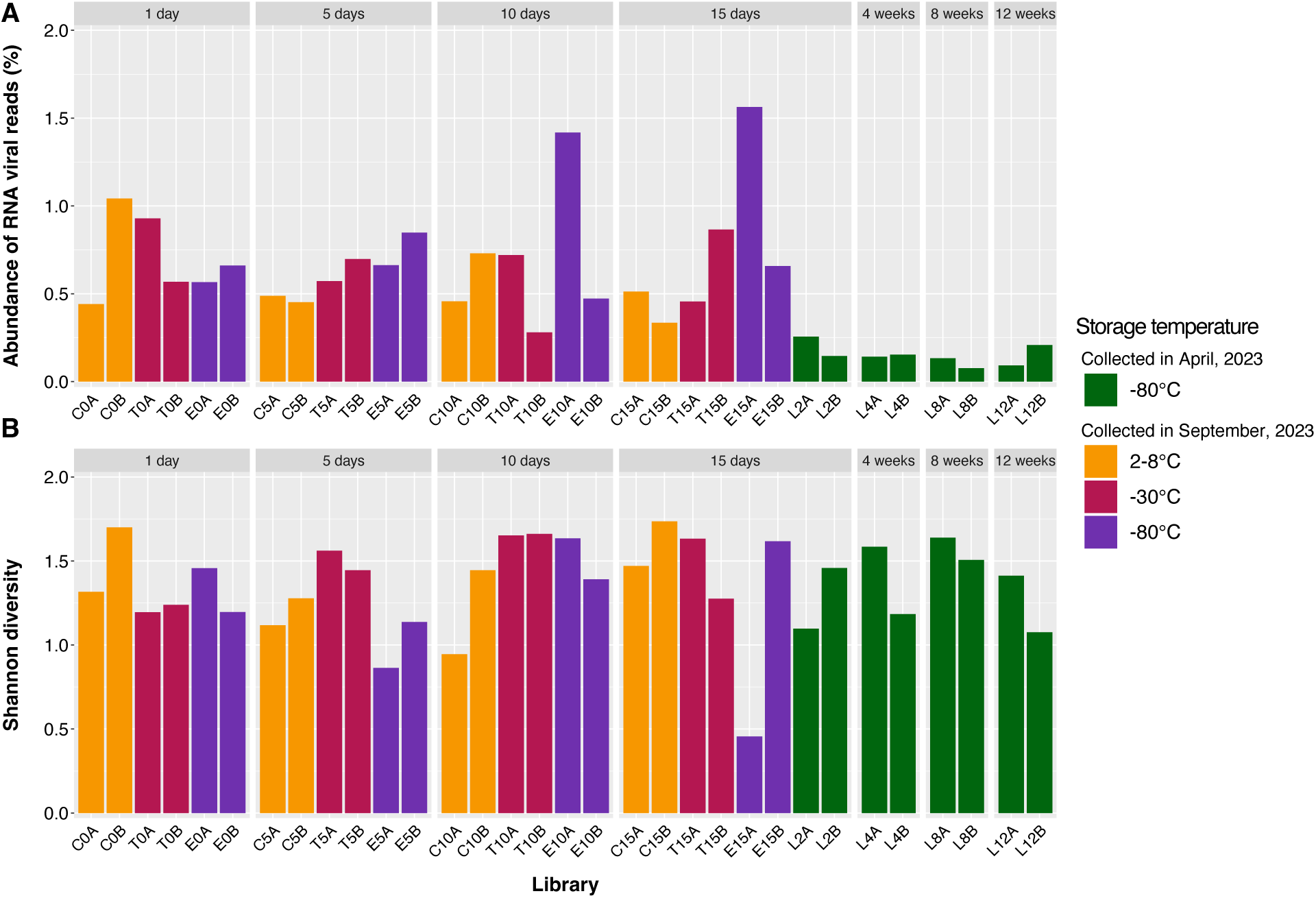
(A) Abundance of reads aligning to RNA virus sequences as a proportion of the total number of reads and (B) Shannon diversity indices of each metatranscriptomic sequence library generated from the Camden soil samples (data sets 1 and 2). Samples stored for up to two weeks at 2-8°C, −30°C, or −80°C are coloured orange, red, and purple, respectively, while samples stored from two to 12 weeks at −80°C are coloured green. Libraries are further grouped by the length of time between collection and extraction (1 day to 12 weeks).

### Effect of storage temperature and time on RNA virome composition

Viral read abundance and alpha diversity indices were compared across the different storage temperatures and lengths of time between collection and extraction to determine their effect on virome composition. Although the samples stored at −80°C for two to 12 weeks (data set 1) had significantly lower viral read abundance (Fig. 2A) and richness (Fig. 2B), these samples were collected at a different time and therefore could not be directly compared to those stored for up to two weeks (data set 2). In addition, there was no decrease in viral read abundance or alpha diversity over the 12 weeks these samples were stored (Fig. 2). Hence, any differences in virome composition between data set 1 and 2 were attributed to the difference in original sample composition at the time of collection. There was also no observable difference in viral read abundance (Fig. 2A) or richness (Fig. 2B) across data set 2, and no significant difference in Shannon diversity indices between any libraries (Fig. 2C, D). These results suggest soil samples remain intact at least three months after sampling when stored at −80°C, as well as remaining stable at refrigeration temperatures (2-8°C) or colder for at least two weeks, without requiring the addition of preservative solutions.

**Figure 2.**
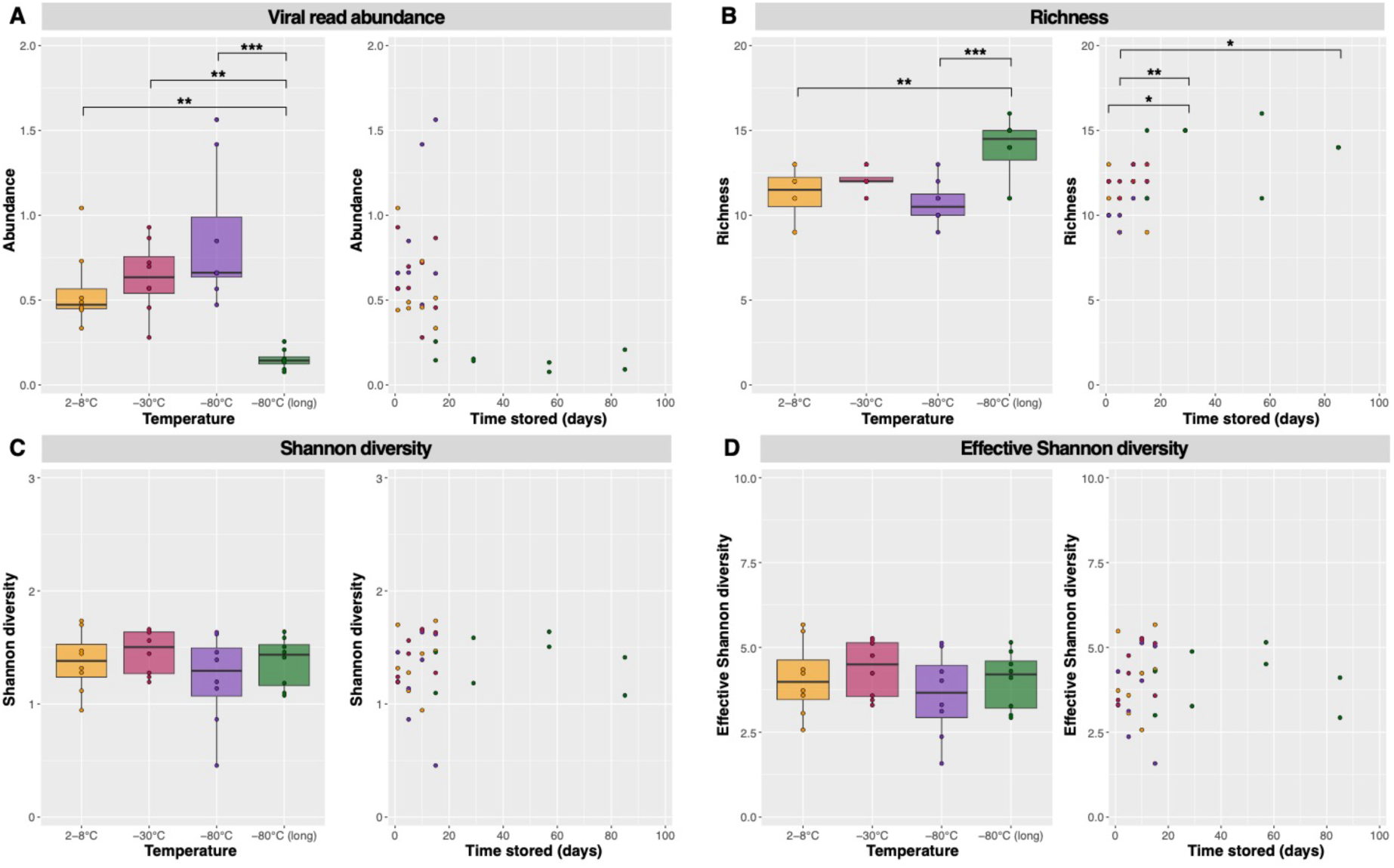
Viral read abundance (A), richness (B), Shannon diversity (C), and effective Shannon diversity (D) indices plotted against storage temperature and length of time stored prior to extraction. Factors significantly different to each other as determined by post-hoc Tukey tests are marked by brackets with one, two, or three asterisks (*) representing p-values less than 0.05, 0.01, and 0.005, respectively.

Overall, members of the RNA virus phylum *Lenarviricota* comprised the vast majority (85%) of viral reads. Specifically, the bacteriophage class *Leviviricetes* within the *Lenarviricota* dominated the soil viromes generated, comprising 57% of viral reads, while the predominantly microbe-associated families *Narnaviridae*, *Mitoviridae*, *Botourmiaviridae*, *Partitiviridae*, and *Picobirnaviridae* were also present in each library, albeit usually at lower proportions (Fig. 3A).

**Figure 3.**
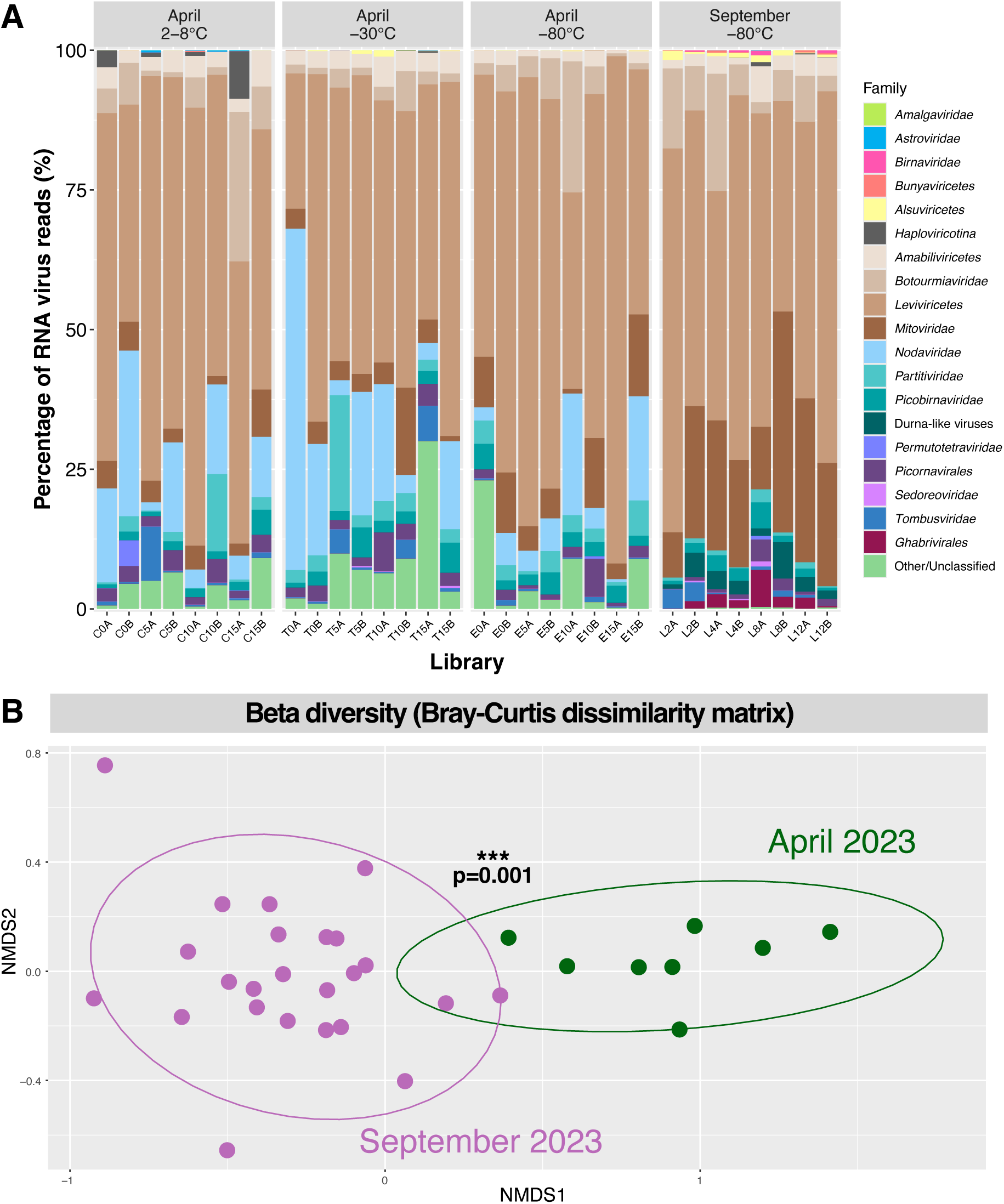
(A) Virome composition at the level of family, order, class, or subphylum of the 32 Camden soil libraries generated in this study. Relative viral read abundance is represented by the reads mapping to each respective RNA viral taxon as a percentage of the total number of reads mapping to *Riboviria* sequences in that library. Libraries are categorised by month of collection (April and September 2023) and the storage temperature prior to RNA extraction (2-8°C, −30°C, and −80°C). (B) Non-metric multidimensional scaling (NMDS) plot of community composition based on the Bray-Curtis dissimilarity matrix between samples comprising data set 1 (April 2023, in green) and data set 2 (September 2023, in purple). Proximity between data points represent similar community compositions and 95% confidence ellipses were drawn, with the asterisks indicating the two communities are significantly different as assessed by PERMANOVA (p = 0.001).

There were notable differences in virome composition between data set 1 and 2, likely due to these samples being collected at two different time points (April and September of 2023) and slightly different locations within Camden. There was a higher proportion of the *Nodaviridae* and *Partitiviridae* in soils from data set 2, with the former not detectable in data set 1. There were also more hits to unclassified viral sequences in data set 2. Conversely, data set 1 had higher proportions of *Mitoviridae*, *Ghabrivirales*, and *Durnavirales*-like sequences (i.e., those that did not fall into any established *Durnavirales* family). In particular, reads mapping to *Durnavirales*-like and *Ghabrivirales* sequences were absent from all but one and two libraries from data set 2 (T5A and T5B/T15A), respectively (Fig. 3A). To investigate these differences in virome composition further, we constructed a Bray-Curtis dissimilarity matrix and visualised it using non-metric multidimensional scaling (NMDS) (Fig. 3B). This revealed a clear distinction between data sets 1 and 2, supported by a PERMANOVA test (p = 0.001). Further testing for the homogeneity of variances revealed no significant difference in variability within each group (p = 0.34), confirming that virome compositions were significantly affected by the two different collection times.

### Identification and phylogenetic analysis of novel viruses

Despite successful soil virome sequencing only taking place at one site on two occasions, a total of 1,475 novel virus sequences were identified in this study (Fig. 4). These viruses spanned the five largest RNA virus phyla, including those with positive-stranded, negative-stranded, and double-stranded genomes. While the viromes of environmental samples typically contain a large proportion of the microbe-associated phylum *Lenarviricota*, a remarkable 90% of the novel viruses identified in this study (1,334 sequences) fell within this group.

**Figure 4.**
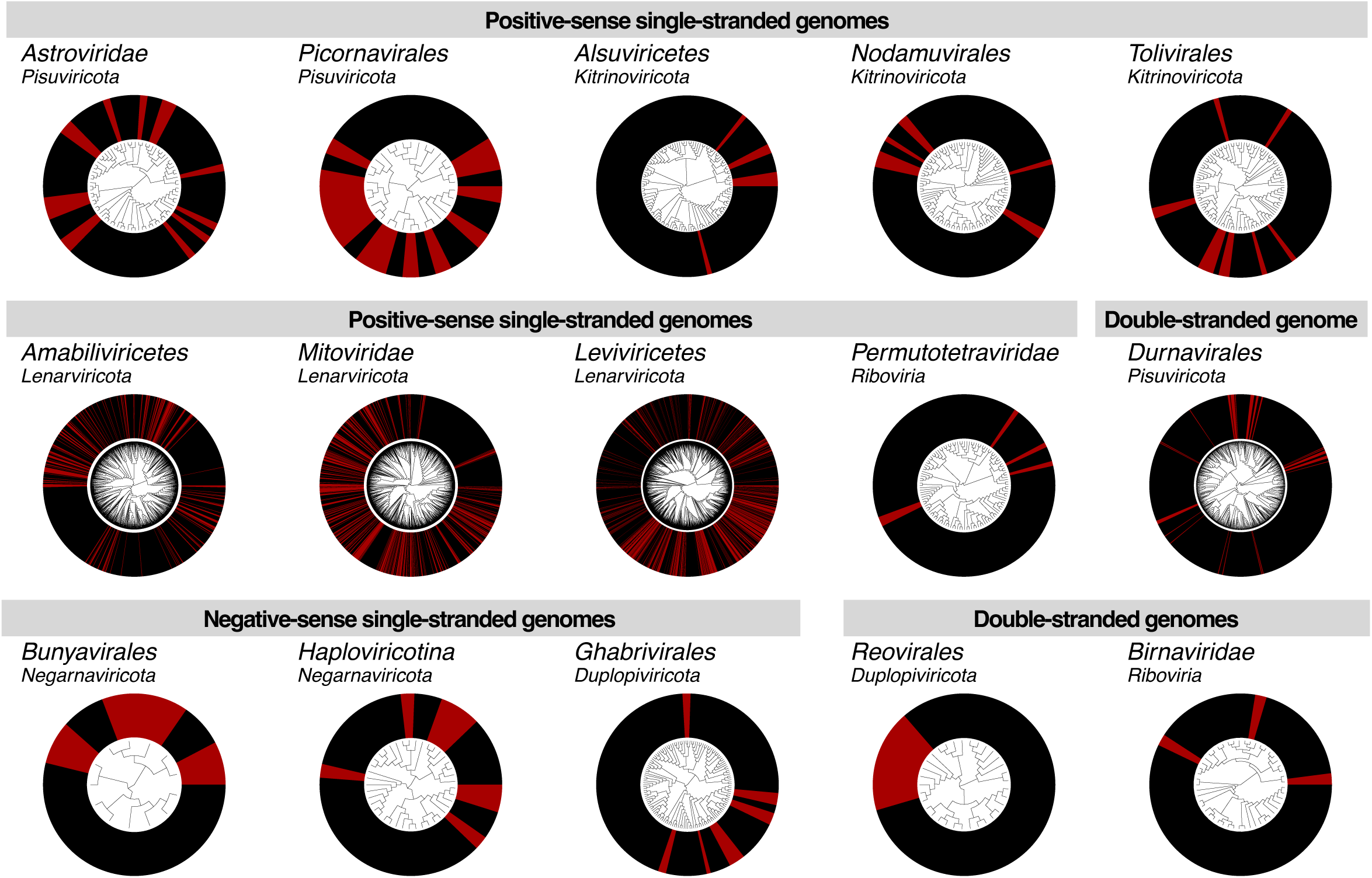
Phylogenetic trees estimated using RdRp amino acid sequences of viruses identified in this study (in red) and related reference sequences (in black). For the purpose of visualisation, all trees are midpoint rooted and phylogenies for certain groups (*Alsuviricetes*, *Amabiliviricetes*, *Durnavirales*, *Ghabrivirales*, *Haploviricotina*, and *Tombusviridae*) were estimated despite containing too much divergence for reliable sequence alignment. Phylogenies for these groups are presented in the supplementary materials and have been divided into multiple, smaller groups containing 1-2 related families each. The sequences in each subset were realigned and trimmed prior phylogenetic inference.

The likely hosts of novel viruses identified in this study were broadly predicted based on the known hosts of related reference sequences. Phylogenetic clustering patterns suggested that viruses with bacterial and fungal hosts comprised 52.4% and 41.9% of novel sequences, respectively, and 1.8% and 1.1% likely had plant and invertebrate hosts, respectively. The remaining 2.8% of sequences were too divergent from reference sequences with a known host to confidently assign a host organism. No vertebrate-associated viruses were identified. Despite the relatively low proportion of plant-associated viruses, these 26 sequences spanned groups with positive-stranded (*Tombusviridae*), negative-stranded (*Qinviridae*, *Yueviridae*), and double-stranded genomes (*Durnavirales*). In contrast, the 772 novel bacteria-associated viruses comprised only two lineages – the class *Leviviricetes* and the recently suggested bacteriophage family *Picobirnaviridae*. Viruses predicted to have invertebrate hosts, of which there were 16, were identified among the *Nodamuvirales*, *Permutotetraviridae*, and *Birnaviridae*, while the 618 likely fungi-associated sequences spanned families within the *Lenarviricota*, *Negarnaviricota*, and *Ghabrivirales*. Notably, 35 of the 42 sequences from the group unable to be assigned host organisms were distantly related to traditionally vertebrate-associated families including the *Astroviridae*, *Picornaviridae*, and *Sedoreoviridae*. The remaining seven had some sequence similarity to plant- and fungi-associated members of the *Alsuviricetes*, but were also too divergent to confidently assign a host.

#### Positive-stranded RNA virus families

The vast majority of the viral diversity identified here fell within the three predominantly positive-sense phyla of RNA viruses – the *Lenarviricota, Pisuviricota*, and *Kitrinoviricota*. We identified novel viruses within each of these phyla, as well as in the family *Permutotetraviridae* that has yet to be assigned a higher taxonomic classification. We now describe each of these groups in turn.

### Lenarviricota

As is frequently observed in environmental samples, the majority of viral diversity observed in our sampling from Camden, NSW, fell into the microbe-associated phylum *Lenarviricota*. Of the 1,334 novel viruses identified in this phylum, 751 belonged to the *Leviviricetes* class of bacteriophage (Fig. S2), and 49, 91, and 380 to the families *Narnaviridae*, *Botourmiaviridae*, and *Mitoviridae*, respectively (Fig. S3-S5). The latter three families are largely fungi-associated, but also contain plant-infecting species and have been detected in several metatranscriptomic data sets generated from vertebrate and invertebrate samples. The remaining 63 viruses fell into a clade of narna-like viruses that has not yet been formally classified but informally referred to as the *Narliviridae* (27) (Fig. S6). Several novel viruses in this phylum were identified in both data sets 1 and 2. Specifically, 29 leviviruses, four narnaviruses, six botourmiaviruses, 14 mitoviruses, and five “narliviruses” were detected in samples collected across both April and September 2023 (Fig. S2-S6).

There was a noticeable difference in the level of phylogenetic novelty between the two major *Mitoviridae* lineages. In the first clade, approximately one-third of sequences were novel (371 of 1116). The second clade was smaller with 318 sequences, of which only nine were novel (∼3%) (Fig. S5). A similar disparity was observed in the *Narnaviridae*, with only five novel narnaviruses present in one lineage of 163 sequences (which also contained the two formally ratified *Narnaviridae* species), and the remaining 44 falling across two lineages totalling 121 sequences (Fig. S3). This indicated that some viral lineages infect hosts that are abundant in Camden soils, while others depend on specific ecological conditions that are not present in every sample. In contrast, novel viruses were present in every genus from the *Botourmiaviridae* (Fig. S4), suggesting this family is associated either with a limited range of microbial hosts or hosts that are equally active across the sampled environment.

### Astroviridae

While astroviruses are typically considered enteric pathogens of animals, recent metatranscriptomic studies of environmental samples have substantially expanded the diversity of this family (12, 15, 39). We identified 15 novel astrovirus-like sequences across all four storage temperatures with representatives from each time point. These sequences fell within a highly diverse clade of environmental astroviruses previously identified in Australian cropping and pasture farmland as well as river sediment from Australia and river water from New Zealand (Fig. 5A; Fig. S7).

**Figure 5.**
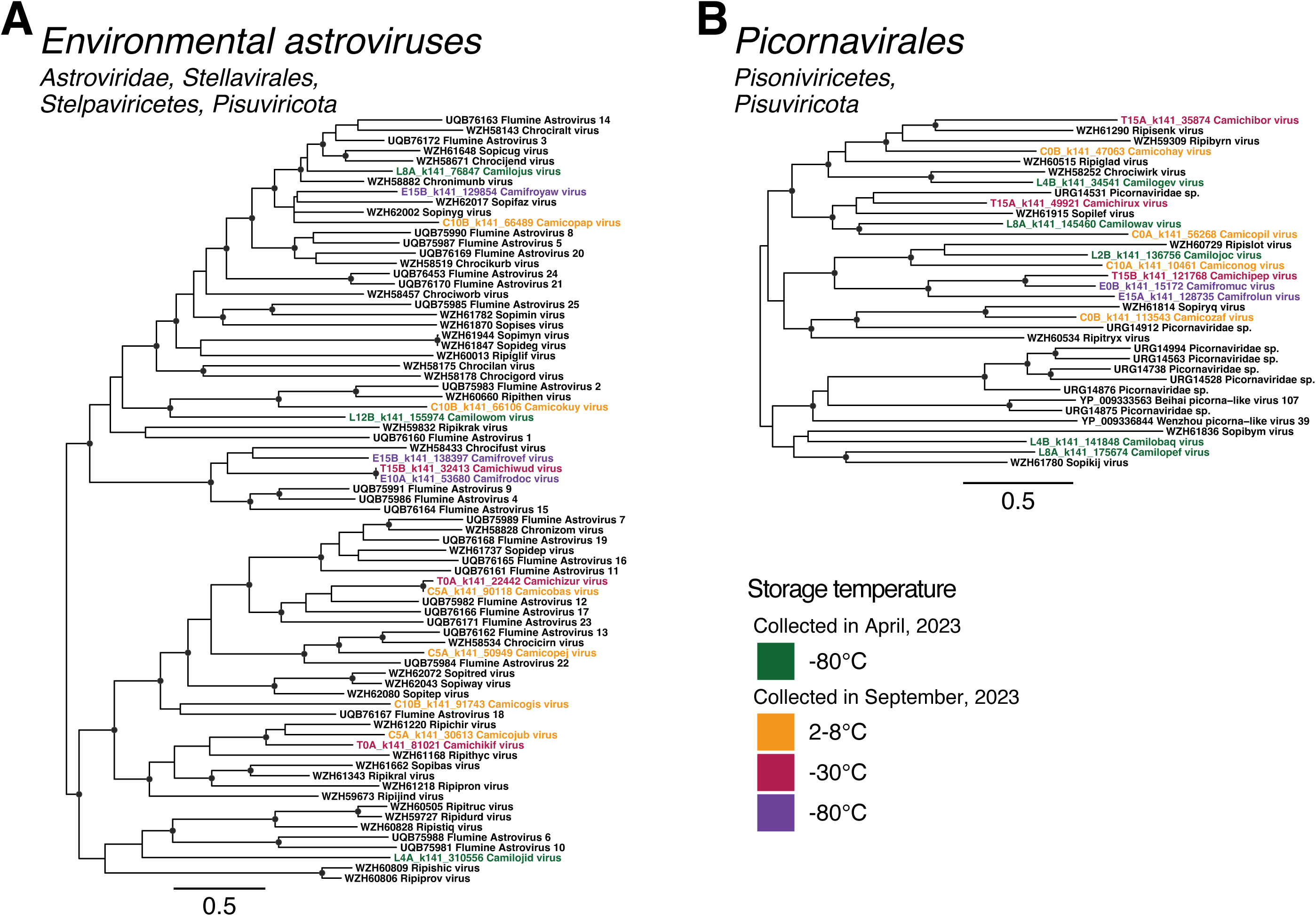
Phylogenetic trees of RdRp amino acid sequences from divergent clades distantly related to the single-stranded, positive-sense RNA virus families (A) *Astroviridae* and (B) *Picornaviridae*. Putative novel viruses identified in this study are coloured by the storage temperature and collection month, such that samples collected in April 2023 (data set 1) and stored at −80°C are coloured in green, while samples collected in September 2023 (data set 2) and stored at 2-8°C are in orange, −30°C in red, and −80°C in purple. Known viruses are coloured in black. Trees are midpoint rooted with branch lengths scaled according to the number of amino acid substitutions per site. Black circles represent node support ≥80% using 1000 SH-aLRT replicates. Individual trees are shown in Fig. S7 and S8.

### Picornavirales

Similar to the *Astroviridae*, all novel picorna-like viruses identified in this study fell within a divergent clade that did not robustly align with members of the *Picornaviridae* associated with vertebrate hosts. Instead, these 14 novel sequences, with representatives found in all four storage temperatures, clustered with divergent picorna-like viruses sourced from Australian and Chinese farmland and sediment samples, as well as Chinese marine invertebrate metagenomes (Fig. 5B; Fig. S8). This clade may represent a new, distinct family within the *Picornavirales*.

### Alsuviricetes

The *Alsuviricetes* is a large class of viruses with a broad host range spanning vertebrates, invertebrates, plants, and fungi. Notable species within this class include alphaviruses, such as Chikungunya virus and Ross River virus, as well as several cropping plant pathogens including tobacco mosaic virus (*Tobamovirus tacabi*), potato virus X (*Potexvirus ecspotati*), and beet necrotic yellow vein virus (*Benyvirus necrobetae*). Despite their host range, there was limited novelty within the *Alsuviricetes* in the data collected here, with only a single divergent sequence identified in the family *Alphaflexiviridae* (Fig. S9A), provisionally named Camofronuj virus (E10B_k141_32818). Based on the midpoint-rooted phylogeny, this virus formed a sister lineage to the plant- and fungi-associated genera *Allexvirus*, *Botrexvirus*, and *Platypuvirus*. An additional six novel sequences were identified in data set 2, falling within a clade of Australian riverbank and pasture farmland samples that was too divergent to robustly align with established families in the *Alsuviricetes* (Fig. S9B). As all 21 sequences comprising this group, including the 15 reference sequences, were obtained from Australian samples, this represents a geographically distinct viral clade.

### Nodaviridae

A total of nine novel noda-like viruses were detected in the samples refrigerated at 2-8°C and frozen at −80°C for up to two weeks (Fig. S10). Two of these fell within relatives to the genus *Alphanodavirus*, with 70.5% and 80.2% amino acid identity to Hainan forest noda-like virus (QYF49887). A single, highly divergent betanoda-like virus was also identified. The remaining six sequences clustered within a third clade predominantly comprising environmental noda-like viruses, but which also contained the alphanodavirus Pariacoto virus. While this clade fell as a sister clade to the betanodaviruses, this node received relatively low statistical support (Fig. S10). As such, the position of this clade cannot be robustly determined; however, this topology has been observed previously (15), suggesting the presence of a third genus within the *Nodaviridae*.

### Tombusviridae

Fewer novel tombusviruses were identified as expected based on previous soil virome studies (11, 12, 15), with only 11 sequences identified across two distinct groups. First, two divergent sequences fell outside of the established members of the subfamily *Regressovirinae* (Fig. S11A). An identical sequence to Camocoruz virus (C0B_k141_18012) was also detected in data set 1 (sequence L2B_k141_345471). Another two novel sequences fell within a group comprised entirely of tombus-like viruses identified in environmental samples (Fig. S5B). Finally, seven novel sequences clustered with a second group of environmental tombus-like viruses that also contained Caledonia beadlet anemone tombus-like virus 1 (ASM94000) (Fig. S11C).

### Permutotetraviridae

Despite the presence of hundreds of permutotetra-like RdRp sequences on NCBI/GenBank derived from a wide variety of sources, including non-Lepidopteran insects, small mammal metagenomes, and environmental samples, the *Permutotetraviridae* currently comprise only two formally ratified species with insect hosts. Herein, four permutotetraviruses were identified from refrigerated samples, while a fifth was identified in soil frozen at −30°C. All five sequences were most closely related to permutotetraviruses previously identified in Chinese soils and sediments (Fig. S12).

#### Negative-stranded RNA virus families

All currently known negative-stranded RNA viruses fall within the phylum *Negarnaviricota*, which comprises two subphyla – the *Haploviricotina* and *Polyploviricotina*. In the case of the subphylum *Haploviricotina*, we identified novel viruses spanning the families *Aspiviridae*, *Qinviridae*, and *Yueviridae* (Fig. S13, 14). Four aspi-like viruses were detected in two distinct clades. The first three robustly grouped with the clade of formally ratified ophioviruses (the single established genus within the family *Aspiviridae*), although Camecokiv virus (C0B_k141_98208) fell outside this clade (Fig. 6A). The fourth sequence – Camecojek (C0B_k141_17830) – was more divergent, with only 30.8% amino acid identity to its closest relative, Erysiphe necator associated negative-stranded RNA virus 12 (QJW70351). This virus fell in the second clade of ophio-like viruses, which may represent a second genus in the *Aspiviridae*, albeit with low node support (Fig. 6A). A single qin-like sequence, provisionally named Camecomuf virus (C10B_k141_5610), was identified and exhibited 44.2% amino acid identity to Qingdao RNA virus 4. Finally, three sequences with sequence similarity (41.5-42.3%) to Goldenrod fern yue-like virus were identified, two of which were detected in the refrigerated samples 15 days post-collection and the third in the sample frozen at −80°C for 12 weeks (Fig. 6B).

**Figure 6.**
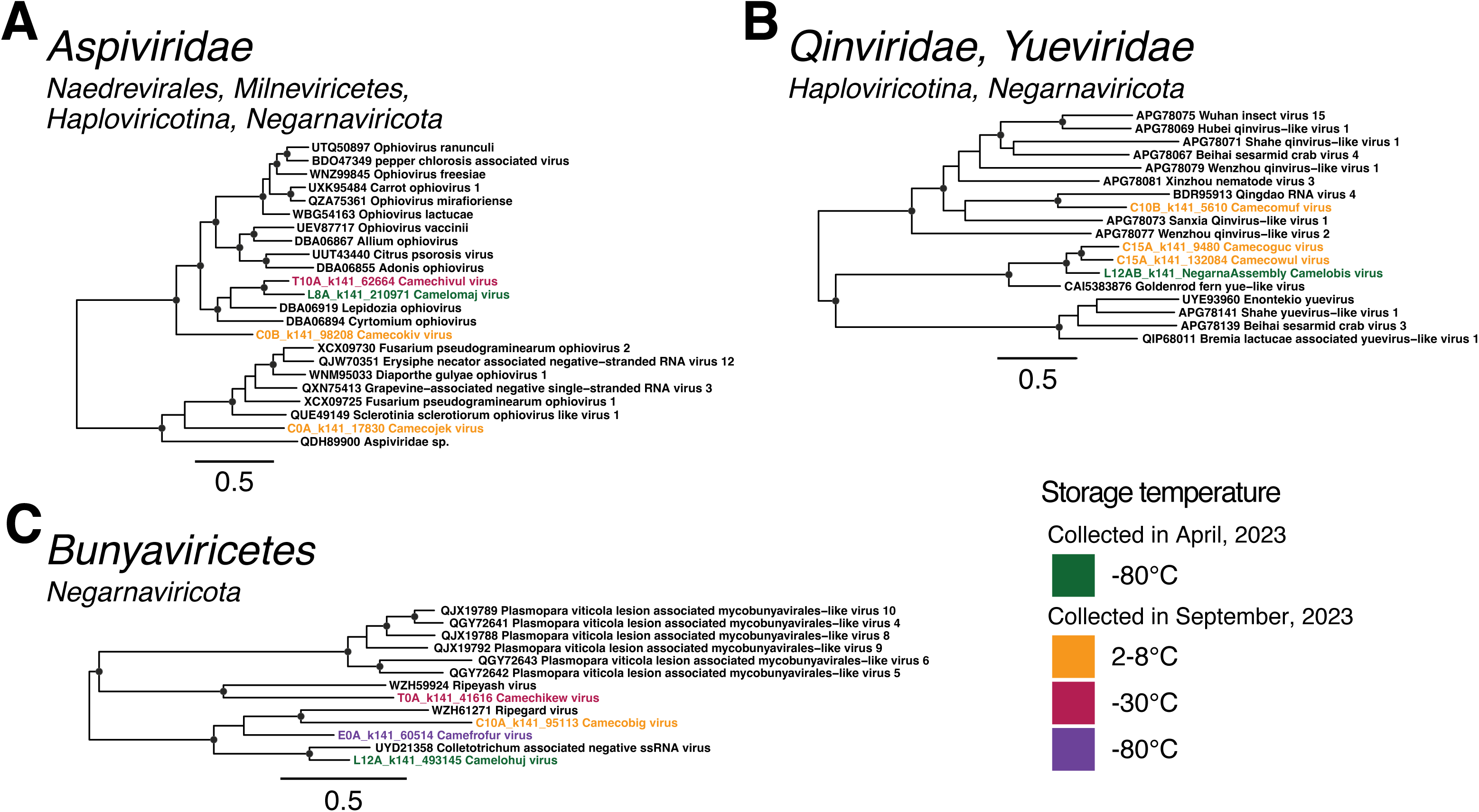
Phylogenetic trees of RdRp amino acid sequences from the single-stranded, negative-sense RNA virus families (A) *Aspiviridae*, (B) *Qinviridae* and *Yueviridae*, and (C) class *Bunyaviricetes*. Putative novel viruses identified in this study are coloured by the storage temperature and collection month, such that samples collected in April 2023 (data set 1) and stored at −80°C are coloured in green, while samples collected in September 2023 (data set 2) and stored at 2-8°C are in orange, −30°C in red, and −80°C in purple. Known viruses are coloured in black. Trees are midpoint rooted with branch lengths scaled according to the number of amino acid substitutions per site. Black circles represent node support ≥80% using 1000 S H-aLRT replicates. Individual trees are shown in Fig. S13-S15.

The subphylum *Polyploviricotina* contains the orders *Bunyaviricetes* and *Articulavirales*, with the former having largely arthropod, plan, protozoan, and vertebrate hosts, while the latter infects predominantly animals. In this study, all novel viruses from this subphylum fell within the *Bunyaviricetes* (Fig. 6C; Fig. S15). Specifically, one sequence from each storage temperature was identified, for a total of four novel bunya-like viruses. However, these did not fall into any established families, instead grouping with the divergent viruses Ripeyash virus (WZH59924), Ripegard virus (WZH61271), and Collectrotrichum associated negative ssRNA virus (UYD21358), which are distantly related to Plasmopara viticola lesion associated mycobunyavirales-like viruses 4-6 and 8-10 (QGY72641-3 and QJX19788, 89, and 92), suggesting these novel viruses may have fungal hosts (Fig. 6C; Fig. S15).

#### Double-stranded RNA virus families

Double-stranded viruses are classified into two RNA viral phyla – *Duplopiviricota* and *Pisuviricota* – the latter of which also contains viruses with positive-sense RNA genomes. Herein, we identified 68 dsRNA viruses across the orders *Durnavirales* (phylum *Pisuviricota*), *Ghabrivirales* and *Reovirales* (phylum *Duplopiviricota*), and the unclassified family *Birnaviridae*.

### Durnavirales

The *Durnavirales* are an order of dsRNA viruses with a host range predominantly comprising fungi, protozoa, and plants, although the *Picobirnaviridae* have recently been proposed to be a family of bacteriophage (28, 30, 40). We identified novel viruses aligning to members of the *Amalgaviridae, Partitiviridae*, and *Picobirnaviridae*, as well as a small group of durna-like viruses that fell outside of any established families (Fig. S16-19).

A single amalgavirus sequence was detected in the −30°C sample extracted 10 days post-collection. This divergent sequence, provisionally named Camichidos virus (T10B_k141_143390), was most closely related (54.6% amino acid identity) to the zybavirus Zygosaccharomyces bailii virus Z (Fig. S16). Second, two alphapartitivirus-like sequences and one betapartitivirus-like sequence were identified. These clustered with previously identified soil and sediment partiti-like viruses and fungi-associated partitiviruses (Fig. S17A, B). From the remaining 11 novel partiti-like viruses, Camicopux virus (C15A_k141_59250) clustered with the deltapartitiviruses and the other 10 fell outside of established genera (Fig. S17C). Five of these formed a distant sister clade to the traditionally protist-associated genus *Cryspovirus*. All four storage temperatures were represented in this sister clade. The *Picobirnaviridae* is a large family with several thousand non-redundant sequences present on GenBank. However, the novel picobirnaviruses identified in this study were restricted to a small subset of sequences, representing the informally proposed genera *Gammapicobirnavirus* and *Etapicobirnavirus* (30). Both proposed genera largely comprise environmentally-sourced picobirnaviruses for which the host organism is uncertain. Two of the novel viruses identified in this study clustered with a group of ten Australian riverbank sediment viruses, resulting in a clade of 12 Australia-specific environmental viruses that form a sister clade to the proposed genus *Etapicobirnavirus*. The remaining 19 novel sequences grouped with the gammapicobirnaviruses. All storage temperatures were represented in the novel picobirnaviruses (Fig. S18). Within the novel sequences identified in the *Durnavirales*, four partiti-like viruses and five picobirna-like vruses were detected across both data sets (Fig. S17C, S18).

Finally, Ulva durnaviruses 1-3 (WIR83939-41) are a group of three viruses previously identified in the metagenome of sea lettuce (*Ulva fenestrata*). These viruses are too divergent to be assigned to a family within the *Durnavirales*. Despite the aquatic nature of this host, eight related viruses were identified in the farmland soil sampled in study, although they were highly divergent with only 29.2-43% amino acid identity to Ulva durnavirus 1 (Fig. 7A; Fig. S19).

**Figure 7.**
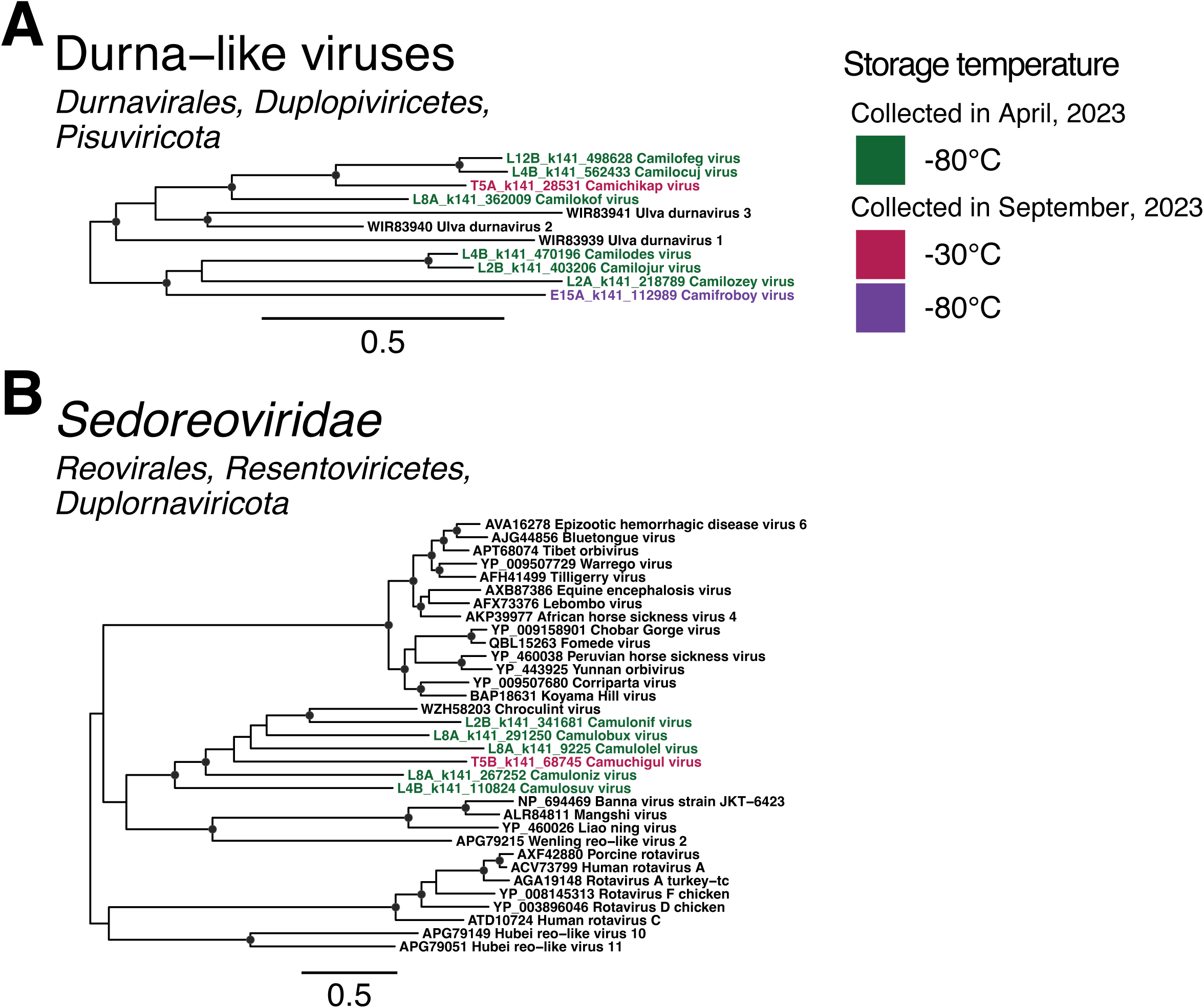
Phylogenetic trees of RdRp amino acid sequences from two double-stranded RNA groups: (A) a divergent clade of unclassified *Durnavirales*-like sequences and (B) the family *Sedoreoviridae*. Putative novel viruses identified in this study are coloured by the storage temperature and collection month, such that samples collected in April 2023 (data set 1) and stored at −80°C are coloured in green, while samples collected in September 2023 (data set 2) and stored at 2-8°C are in orange, −30°C in red, and −80°C in purple. Known viruses are coloured in black. Trees are midpoint rooted with branch lengths scaled according to the number of amino acid substitutions per site. Black circles represent node support ≥80% using 1000 S H-aLRT replicates. Individual trees are shown in Fig. S18 and S20.

### Ghabrivirales

We identified 15 sequences with similarity to members of the *Totiviridae*. Two of these clustered with a group of environmental toti-like viruses that fell outside of the established families *Chrysoviridae* and *Megabirnaviridae*. The remaining 13 fell in two toti-like clades that did not robustly align with any formally ratified families (Fig. S20). The vast majority of reference sequences in this group were identified in soil and sediment samples, potentially representing a distinct family within the order. All but one novel virus in this order were identified in soil comprising data set 1 (stored at −80°C), with the remainder present in a sample from data set 2 that was stored at −30°C for 15 days.

### Reovirales

Novel sequences related to those from the family *Sedoreoviridae* were also identified, five of which were detected in samples stored at −80°C for two to eight weeks, and the sixth in a sample stored at −30°C. These viruses were most closely related to an Australian cropping farmland virus, Chroculint virus (WZH58203). However, these divergent sequences did not cluster within any established genera. Instead, they formed a sister clade to a group containing Banna virus (NP_694469), Mangshi virus (ALR84811), Liao ning virus (YP_460026), and Wenling-reo-like virus (APG79215) (Fig. 7B; Fig. S21).

### Birnaviridae

The family *Birnaviridae* is yet to be assigned to a taxonomic level higher than family. This family is associated with a wide range of hosts including birds, fish, and insects, to more divergent organisms such as rotifers and molluscs. These viruses largely group by host organism on phylogenetic trees, although bird- and fish-associated genera typically form a monophyletic group, and insect-associated birnaviruses comprise two, phylogenetically distinct genera. Aquatic environments such as sediment and river water have also yielded a large number of birna-like viruses that fall outside the established genera. In this study, three birna-like sequences were identified in samples stored at −80°C for four to eight weeks, clustering with a group of exclusively environmentally-sourced sequences that formed a sister clade to formally ratified *Birnaviridae* species (Fig. S22). These novel birna-like viruses were also divergent, with 22.9-30% amino acid identity to their closest relatives, Xinjiang sediment birna-like virus 1 (UMO75744), Zhejiang birna-like virus (UMO75762), and Yunnan birna-like virus 2 (UMO75742).

## Discussion

We identified 1,475 novel RNA virus sequences from the 32 libraries generated in this study. Novel viruses were present in the five largest RNA viral phyla, with the vast majority (90%) of diversity present in the microbe-associated phylum *Lenarviricota*, as is typical of terrestrial environments (11, 12, 15, 28). Notably, viruses with sequence similarity to the *Astroviridae* and *Picornaviridae* – families traditionally thought to be animal-infecting – formed divergent, monophyletic clusters with other environmentally-sourced sequences, potentially representing distinct new families and possibly expanding the host ranges of these groups to invertebrates or fungi, which are abundant in soil systems. As all 32 libraries were generated from soil samples taken from a single site (The University of Sydney’s Camden campus) across two collection time points, this result highlights the remarkably rich viral diversity present in soils in Australia and perhaps globally. Importantly, we observed instances of country-specific evolution in the *Astroviridae*, *Alsuviricetes*, *Mitoviridae*, “*Narliviridae*”, *Partitiviridae*, *Picobirnaviridae*, *Ghabrivirales*, and *Sedoreoviridae*, where novel viruses formed clusters with previously published sequences derived exclusively from Australian (and New Zealand, in the case of the *Astroviridae*) environmental data sets.

Within the families *Narnaviridae* and *Mitoviridae* we also observed lineages with a disproportionately low number of novel viruses compared to other lineages from the same virus families. This is unlikely to be driven by differences in sample type or geographical location as soil- and sediment-derived sequences were evenly distributed across both phylogenies. While this could be due to sufficient sampling of certain environments, it seems more likely that these viruses are underrepresented in Camden soil and depend on specific hosts or ecological conditions not met here. In contrast, the viruses in the high-novelty lineages may have more abundant or active microbial hosts in the sampled soils. These results highlight the importance of sampling an extensive range of terrestrial environments to uncover divergent viral lineages that may be restricted to specific ecological and geographical conditions, or infect hosts occupying a limited range of environmental niches.

Approximately half of the novel viruses identified (751 out of 1,475) represented the bacteriophage order *Leviviricetes*, with the families *Narnaviridae*, *Mitoviridae*, *Botourmiaviridae*, *Partitiviridae*, and *Picobirnaviridae* also universally present. From these groups a subset of 40 novel viruses, along with a novel tombusvirus, were detected in both data set 1 and data set 2, despite their collection taking place in April (autumn) and September (spring) of the same year, respectively, and from slightly different locations in Camden. However, there were also interesting differences between the two data sets, including the presence of some groups, such as the *Nodaviridae*, *Ghabrivirales*, and durna-like viruses at one collection time that were almost, if not entirely, absent from the other. These differences in virome composition between data sets (i.e., the beta diversity of these samples) were statistically significant. While it is not uncommon for virome compositions to vary across soil types and land uses (12, 13), it is notable for such distinct differences to occur within a single site. These data showcase the remarkable temporal and spatial variability of terrestrial environments and highlight the necessity to take multiple replicates per site and generate comprehensive ecological metadata to produce accurate representations of soil viral communities (41, 42).

Despite the impressive diversity present in these samples, we found no statistically significant difference in viral abundance or diversity among samples that were collected simultaneously, regardless of the storage condition. Interestingly, we did observe variation in virome composition among samples collected from the same overall location but five months apart, suggesting that temporal dynamics and soil heterogeneity, rather than storage-related degradation, may play a larger role in shaping soil viral communities. This aligns with previous findings that soil viromes are highly diverse and dynamic, reflecting both microbial turnover and environmental fluctuations (43, 44). While we had hypothesised that warmer storage temperatures and prolonged delays would degrade RNA quality or bias sequencing outputs, our statistical analyses revealed no measurable effect of storage time or temperature on RNA yields, integrity, sequencing success, or virome composition. This is a promising result for researchers conducting fieldwork in remote locations, where immediate access to ultra-cold storage may not be feasible.

Surprisingly, the three commercial RNA preservative solutions tested (over multiple sampling time points and locations) were consistently less effective than simply freezing soil samples without any additive. Hence, the heterogeneous, reactive nature of soil is likely incompatible with targeted chemical solutions. On a chemical level, organic matter and soil minerals such as clay and metal oxides may present barriers through their interactions with both RNA and preservative solutions (34, 45). For example, RNA binds to these soil particles, preventing preservatives from accessing and forming protective complexes with RNA as it would in tissue samples. RNA adsorption to soil particles is enhanced by the presence of divalent cations such as Ca^2+^ and Mg^2+^ (34), further reducing accessibility by a preservative solution. Hence, when applied to soil, chemical preservative solutions are outperformed by the immediate and uniform slowing of enzymatic activity accomplished by snap freezing or refrigerating samples soon after collection.

We further note that we successfully generated metatranscriptomic data from a single, managed farmland site (Camden, NSW). Agricultural interventions and human activity can have a profound impact on microbial abundance and diversity. Generally, crop monocultures, intensive fertilisation, irrigation, and tillage can contribute to soil microhabitat destruction and reduced nutrient availability, resulting in decreased microbial host abundance (46, 47). Conversely, certain interventions have been observed to increase microbial abundance and diversity, such as the application of phosphorus-based fertilisers that can be utilised by fungal hosts for growth, or mixing via tillage that can increase bacterial diversity by reducing the spatial separation of microbes and key substrates needed for replication (48, 49). Hence, virome composition is likely to vary significantly even across differently managed agricultural lands. Soils from other biomes, such as urban, forested, or pristine environments, or sediment adjacent to aquatic and marine environments, will harbour even more distinct microbial communities and virome compositions compared to farmland soils (13, 47, 50). It is therefore plausible that viral stability under the same conditions as this study may vary across soils from different ecological systems not tested here. Moreover, our experimental design lacked a true ambient-temperature control (i.e., a sample stored at room temperature without intervention), which limits direct comparison to a potential “worst-case” field scenario. While this reduces the universal applicability of our results, these data still provide practical guidelines for the fieldwork elements of viral metatranscriptomic studies, including reasonable expectations for sample handling timelines and minimum storage conditions that do not compromise data quality.

The detection of 1,475 novel RNA viruses from a single sampling site corroborates recent literature highlighting the remarkable, untapped viral diversity in terrestrial environments, particularly in Australian soils. Noticeable differences in virome compositions between samples collected from the same site five months apart show that the spatiotemporally dynamic nature of soil is reflected in its viruses. Importantly, these data show that short-to mid-term storage under refrigeration or freezing conditions does not result in a detectable loss in viral abundance, diversity, or sequencing quality. This has positive implications for the accessibility of remote soil sites, which likely harbour even greater viral diversity. Sampling these sites and collecting extensive associated physiochemical metadata will result in substantial expansions of the RNA virosphere and further reveal the complex relationships between environmental viruses and the intricate soil networks surrounding them.

## Materials and Methods

### Sample collection and storage conditions

All soil was collected from sites in Sydney, New South Wales, Australia, conducted across four rounds of sampling. The first two rounds (September 2022 and February 2023) took place in a residential garden in Marrickville, NSW. This site featured a brown loamy topsoil with no fertilisation or tilling history within the three years prior to sample collection.

(1) In September 2022, a total of 7g of soil was collected into six tubes each containing 28mL of PowerProtect DNA/RNA (Qiagen), then transported to laboratory facilities at room temperature. Three tubes were stored at room temperature, while the remaining three tubes were moved to −80°C freezer for one week, at which point RNA extraction was performed.
(2) In February 2023, 2g of soil was collected into tubes containing 2mL of DNA/RNA Shield (Zymo Research), 4mL of DNA/RNA Shield, and 18mL of RNA*later* (Thermo Fisher Scientific), with four tubes per preservative solution. These samples were transported to laboratory facilities at room temperature, with two tubes from each set then moved to refrigeration at 2-8°C until RNA extraction three days later. The remaining samples were stored at room temperature for three days, then moved to 2-8°C for three days, and RNA was extracted a total of six days after collection. The next two rounds (April 2023 and September 2023) were conducted at The University of Sydney’s Camden campus from the same paddy site (Lat. −34.02, Long. 150.66) of fallow land. This site harboured a loamy sand soil surface horizon with a pH of 6.2 (measured using a 1:5 soil:water suspension test) and a total soil carbon content of approximately 1%.
(3) In April 2023, 2g of soil was collected into 40 tubes containing 2mL DNA/RNA Shield each. These samples were transported to laboratory facilities at room temperature. Upon arriving, 20 samples were placed in refrigeration conditions at 2-8°C and the remainder were stored at room temperature. At two, four, eight, and 12 weeks after the date of collection, two samples were moved from room temperature to refrigeration conditions and from refrigeration to frozen at −80°C. RNA extraction was performed on two samples per storage condition at each of these time points to capture the RNA profile of samples over time at various temperature. In addition, during sampling, two 50mL falcon tubes were filled with soil without the addition of any preservative and stored at −80°C immediately upon delivery to laboratory facilities. RNA from these control samples was extracted at the two-, four-, eight-, and 12-week mark alongside the preserved samples.
Finally, (4) in September 2023, six 50mL tubes were filled with soil and transported back to laboratory facilities on ice. No preservative solution was used. Immediately upon returning to the laboratory, two tubes were placed at 2-8°C, −30°C, and −80°C, respectively. RNA was extracted from each sample within 24 hours of collection, then again at five, ten, and 15 days post-collection.

### RNA extraction, library construction and sequencing

For all soil samples, RNA was extracted using the RNeasy PowerSoil Total RNA kit (Qiagen) as per the manufacturer’s instructions with an additional incubation of 10 minutes at room temperature following the addition of phenol-chloroform-isoamyl alcohol (and vortexing briefly to mix), prior to vortexing for 15 minutes. For samples preserved in PowerProtect DNA/RNA or RNA*later*, the preservative solution was removed prior to extraction by centrifuging at 3,000 x *g* for 15 minutes and decanting the supernatant. Samples were then washed by adding and mixing 24mL of PBS, centrifuging at 3,000 x *g* for 30 minutes, then decanting the supernatant. This washing procedure was performed twice before beginning the RNA extraction protocol with 1-2g of soil sample. For samples preserved in DNA/RNA Shield, the entire contents of the tube were used in the extraction due to the lytic nature of DNA/RNA Shield. Extracted RNA was quantified using the Qubit RNA high sensitivity (HS) Assay Kit on the Qubit Fluorometer v3.0 (Thermo Fisher Scientific) and stored at −80°C prior to library construction and sequencing. Libraries were constructed using the Illumina Stranded Total RNA library preparation protocol and rRNA was removed using the Ribo-Zero Plus rRNA depletion kit (Illumina). In cases of successful library construction, libraries were sequenced on the Illumina NovaSeq 6000 platform (paired-end, 150 bp). Library preparation and sequencing was performed by the Australian Genome Research Facility (AGRF).

### Data processing

Bioinformatic analyses were performed on The University of Sydney’s Artemis high-performance computing system. The initial processing of sequencing data utilised the BatchArtemisSRAMiner workflow (51). Sequence reads were adaptor- and quality-trimmed using Trimmomatic (v0.38) (52), then assembled into contigs using MEGAHIT (v1.2.9) (53) with the default assembly parameters. No eukaryotic or bacterial reads were filtered prior to assembly. Sequence quality was checked for both raw reads and trimmed reads using FastQC (v0.11.8) (54). FastQC was also used to confirm the successful removal of adaptors in trimmed reads. Assembled contigs were compared to an in-house curated database of viral sequences using DIAMOND BLASTX (v2.0.9) (55) with an e-value of 1 × 10^−5^ for sensitivity. This database was curated in January 2024 by querying NCBI’s protein sequence database for entries with ‘Viruses’ in the organism field and built using the ‘makedb’ function in DIAMOND BLAST. Contigs returning positive hits were compared using DIAMOND BLASTX to the non-redundant (nr) protein database as of January 2024 to identify false positives. Contigs with hits to viral sequences were retained and sorted by the taxonomy of the closest relative indicated by the DIAMOND BLASTX results.

Contigs over 1000 nucleotides (nt) in length with DIAMOND BLASTX hits to the viral RNA-dependent RNA polymerase (RdRp) were retained for phylogenetic analysis. These contigs were translated to amino acid sequences, generally using the standard genetic code (i.e., translation table 1), aside from the family *Mitoviridae* where using the mitochondrial genetic code (translation table 4) was required to produce uninterrupted ORFs of the expected length. Translated sequences were checked in Geneious Prime 2025.1.3 for the presence of the conserved A, B, and C motifs characteristic of the RNA virus RdRp. Contigs containing these motifs were considered likely to represent RNA viruses and were used in subsequent phylogenetic analyses.

Amino acid sequences from each taxonomic group, including previously established viruses as reference sequences, were aligned using MAFFT (v7.402) (56) and trimmed using TrimAl (v1.4.1) (57) to retain the most conserved positions, producing final multiple sequence alignments of approximately 700 amino acids in length. Phylogenetic trees were then estimated using the maximum likelihood approach in IQ-TREE (v1.6.12) (58) with 1000 SH-aLRT replicates to evaluate node support. The resultant trees were visualised using the ‘ape’ (59) and ‘ggtree’ (60) packages in R. The final set of contigs determined to represent novel RNA viruses by these phylogenetic analyses were then compared once more to the nr protein database as of May 2025 using DIAMOND BLASTX to achieve the most up-to-date records for their closest published relative and are described in Table S1.

### Viral abundance and diversity comparison

The relative abundance of viral transcripts within each sequencing library was estimated using RSEM (v1.3.1) (61). Viral read abundance was calculated as the expected count of reads mapping to viral contigs divided by the total raw read count x 100. Virome compositions for each library were calculated as the expected count of reads aligning to each viral family/order/class/subphylum as a proportion of the total expected count of reads aligning to RNA virus sequences (i.e., *Riboviria*). Higher taxonomic levels were used when certain families comprised too small a proportion of the virome to be adequately visualised (such as the *Alsuviricetes*, *Bunyaviricetes*, *Haploviricotina*, and *Ghabrivirales*), or sequences were too divergent to be assigned at the family level (*Amabiliviricetes*, *Leviviricetes*, and *Picornavirales*).

Viral alpha diversity indices (i.e., the diversity within a sample) were calculated in RStudio (v2025.05.0+496) (62), R (v4.5.1) (63), using an adapted Rhea alpha diversity script (64) as described by Wille *et al*. (2018) (65). Alpha diversity indices include richness, Shannon diversity, and effective Shannon diversity (calculated as the exponential of each respective Shannon diversity index). To include divergent sequences that did not robustly fall in any established families or genera, alpha diversity was calculated at the order level, meaning that for the purposes of this study richness refers to the number of unique orders detected within a library. Along with viral read abundance and alpha diversity, the effect of storage conditions on extraction RNA concentrations (normalised for the input mass of soil used in each extraction), RNA Integrity Number (RIN), and sequencing data yield were also investigated. Beta diversity (i.e., the diversity between samples) was assessed using the Bray-Curtis dissimilarity matrix, calculated using the ‘vegdist’ function, and the homogeneity of group dispersions was assessed using the ‘betadisper’ function, both from the R ‘vegan’ package (66). The effects of storage temperature, time, and month of collection on virome composition were statistically evaluated using the PERMANOVA (Adonis tests), also using the ‘vegan’ package. The dissimilarity between viral community compositions was visualised using non-metric multidimensional scaling (NMDS) with the ‘phyloseq’ package (67).

In all cases, temperature was treated as a categorical variable, such that its effect on viral read abundance and alpha diversity was tested using generalised linear models. The influence of time in storage prior to extraction was tested using linear models. The significance of these models was evaluated using χ^2^ tests and significant differences between pairs of variables were determined using post-hoc Tukey tests. The effects of time and temperature on data set 1 (libraries beginning with “L”) were not directly compared to data set 2 (libraries beginning with “C”, “T”, and “E”), as they represent different original soil samples, therefore introducing many confounding variables that likely affect viral read abundance and alpha diversity.

## Acknowledgements

This work was funded by an Australian Commonwealth Department of Agriculture, Water and the Environment grant (K27RZJZ) to BM, ABM and ECH.

## Data Availability

The raw sequence read data for this study are available in the NCBI Sequence Read Archive (SRA) database under the BioProject PRJXXXX and SRA accession numbers XXX – XXX. All genome sequences used in the phylogenetic analysis are available in NCBI GenBank under the accession numbers XXX – XXX.

