## Supplementary material for "Limited effect of short- to mid-term storage conditions on an Australian farmland soil RNA virome": Figures S1-S2

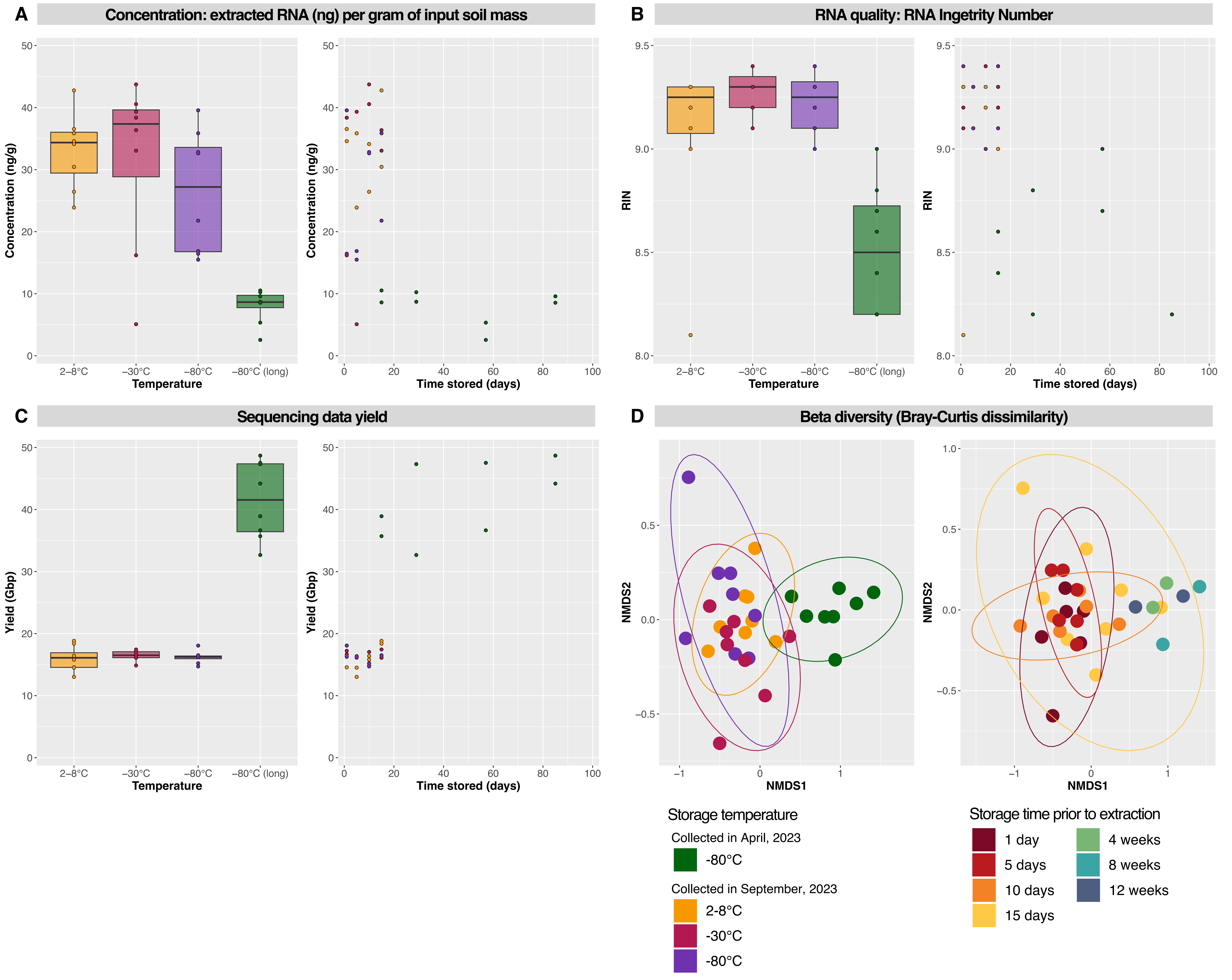

A
Concentration: extracted RNA (ng) per gram of input soil mass
Concentration (ng/g)
Temperature
Time stored (days)
2-8°C
−30°C
−80°C
−80°C (long)
B
RNA quality: RNA Ingetrity Number
RIN
C
Sequencing data yield
Yield (Gbp)
D
Beta diversity (Bray-Curtis dissimilarity)
NMDS1
NMDS2
Storage temperature
Collected in April, 2023
-80°C
Collected in September, 2023
2-8°C
-30°C
-80°C
Storage time prior to extraction
1 day
5 days
10 days
15 days
4 weeks
8 weeks
12 weeks

**Supplementary Figure 1.** Extracted RNA concentration per gram of input soil (A), extracted RNA quality (B), and sequencing data yield (C) plotted against storage temperature and length of time stored prior to extraction. (D) Non-metric multidimensional scaling (NMDS) plot of community composition based on the Bray-Curtis dissimilarity matrix between samples grouped by storage temperature or length of time stored prior to extraction. Proximity between data points represent similar community compositions and 95% confidence ellipses were drawn.

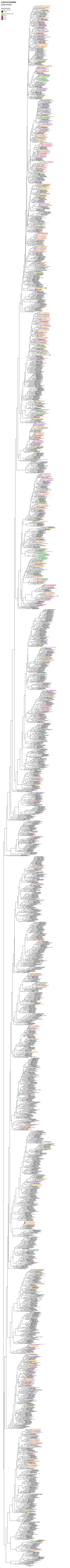
Leviviricetes
Lenarviricota
Storage temperature
Collected in April, 2023
-80°C
Collected in September, 2023
2-8°C
-30°C
-80°C

**Supplementary Figure 2.** Phylogenetic tree of RdRp amino acid sequences from the class *Leviviricetes*. Putative novel viruses identified in this study are coloured by the storage temperature and collection month, such that samples collected in April 2023 (data set 1) and stored at -80°C are coloured in green, while samples collected in September 2023 (data set 2) and stored at 2-8°C are in orange, -30°C in red, and -80°C in purple. Known viruses are coloured in black. Sequences found in both data sets are indicated by a coloured circle next to the provisional virus name, coloured by the storage temperature of library or libraries the identical sequence was found in. Trees are midpoint rooted with branch lengths scaled according to the number of amino acid substitutions per site. Black circles represent node support ≥80% using 1000 S H-aLRT replicates.

### *Narnaviridae*

*Amabiliviricetes, Lenarviricota*

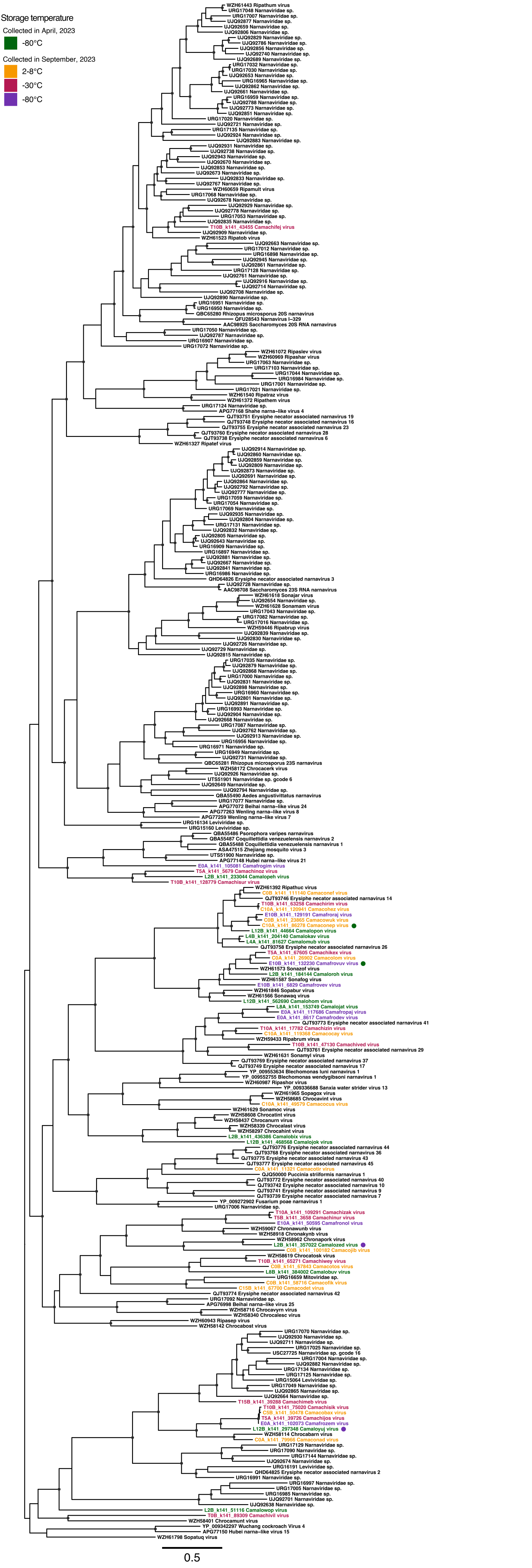

**Supplementary Figure 3.** Phylogenetic tree of RdRp amino acid sequences from the family *Narnaviridae*. Putative novel viruses identified in this study are coloured by the storage temperature and collection month, such that samples collected in April 2023 (data set 1) and stored at -80°C are coloured in green, while samples collected in September 2023 (data set 2) and stored at 2-8°C are in orange, -30°C in red, and -80°C in purple. Known viruses are coloured in black. Sequences found in both data sets are indicated by a coloured circle next to the provisional virus name, coloured by the storage temperature of library or libraries the identical sequence was found in. Trees are midpoint rooted with branch lengths scaled according to the number of amino acid substitutions per site. Black circles represent node support ≥80% using 1000 S H-aLRT replicates.

### *Botourmiaviridae*
#### *Miaviricetes, Lenarviricota*

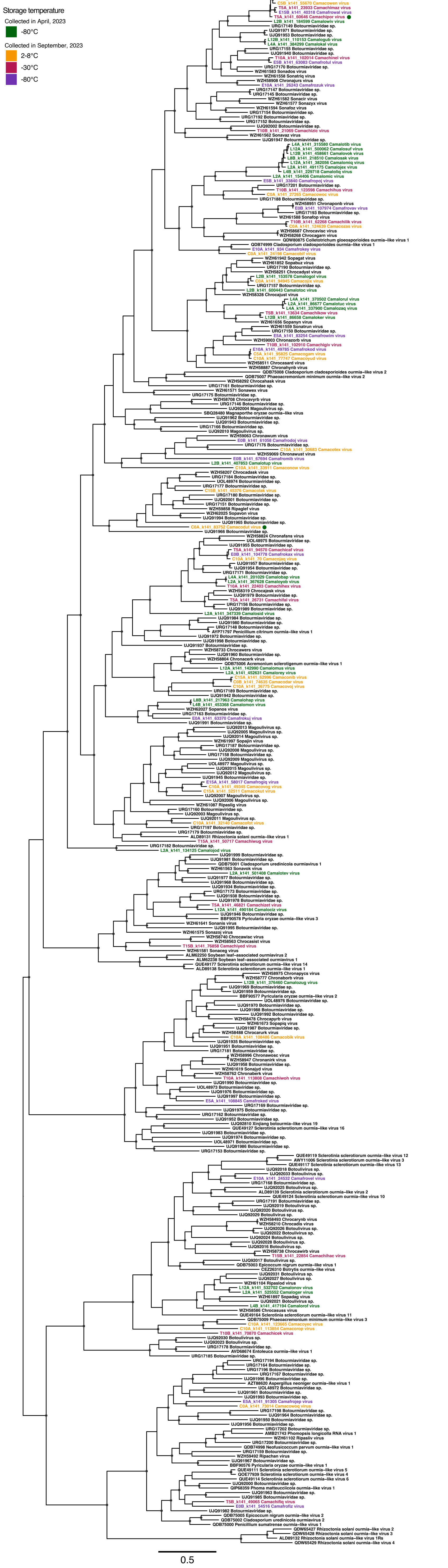

**Supplementary Figure 4.** Phylogenetic tree of RdRp amino acid sequences from the family *Botourmiaviridae*. Putative novel viruses identified in this study are coloured by the storage temperature and collection month, such that samples collected in April 2023 (data set 1) and stored at -80°C are coloured in green, while samples collected in September 2023 (data set 2) and stored at 2-8°C are in orange, -30°C in red, and -80°C in purple. Known viruses are coloured in black. Sequences found in both data sets are indicated by a coloured circle next to the provisional virus name, coloured by the storage temperature of library or libraries the identical sequence was found in. Trees are midpoint rooted with branch lengths scaled according to the number of amino acid substitutions per site. Black circles represent node support ≥80% using 1000 S H-aLRT replicates.

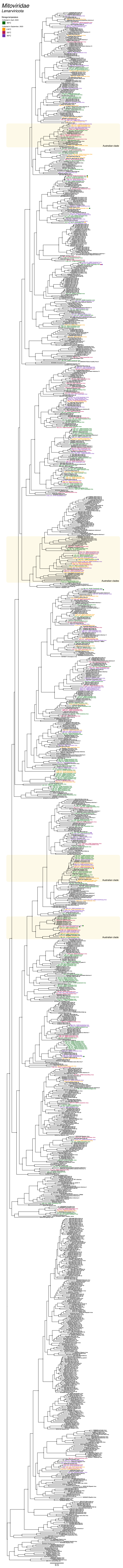
Mitoviridae
Lenarviricota
Storage temperature
Collected in April, 2023
-80°C
Collected in September, 2023
2-8°C
-30°C
-80°C
Australian clade
Australian clades
Australian clade
Australian clade
0.5

**Supplementary Figure 5.** Phylogenetic tree of RdRp amino acid sequences from the family *Mitoviridae*. Putative novel viruses identified in this study are coloured by the storage temperature and collection month, such that samples collected in April 2023 (data set 1) and stored at -80°C are coloured in green, while samples collected in September 2023 (data set 2) and stored at 2-8°C are in orange, -30°C in red, and -80°C in purple. Known viruses are coloured in black. Sequences found in both data sets are indicated by a coloured circle next to the provisional virus name, coloured by the storage temperature of library or libraries the identical sequence was found in. Clades comprised exclusively of sequences sourced from Australian environmental samples are indicated by a yellow box. Trees are midpoint rooted with branch lengths scaled according to the number of amino acid substitutions per site. Black circles represent node support ≥80% using 1000 S H-aLRT replicates.

'Narliviridae'

*Amabiliviricetes, Lenarviricota*

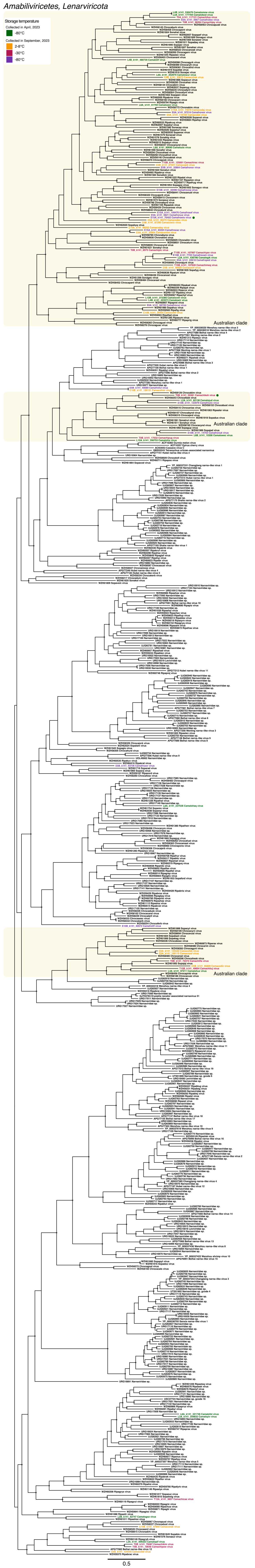

**Supplementary Figure 6.** Phylogenetic tree of RdRp amino acid sequences from the proposed family *'Narliviridae'*. Putative novel viruses identified in this study are coloured by the storage temperature and collection month, such that samples collected in April 2023 (data set 1) and stored at -80°C are coloured in green, while samples collected in September 2023 (data set 2) and stored at 2-8°C are in orange, -30°C in red, and -80°C in purple. Known viruses are coloured in black. Sequences found in both data sets are indicated by a coloured circle next to the provisional virus name, coloured by the storage temperature of library or libraries the identical sequence was found in. Clades comprised exclusively of sequences sourced from Australian environmental samples are indicated by a yellow box. Trees are midpoint rooted with branch lengths scaled according to the number of amino acid substitutions per site. Black circles represent node support ≥80% using 1000 S H-aLRT replicates.

### *Environmental astroviruses*

#### *Astroviridae, Stellavirales, Stelpaviricetes, Pisuviricota*

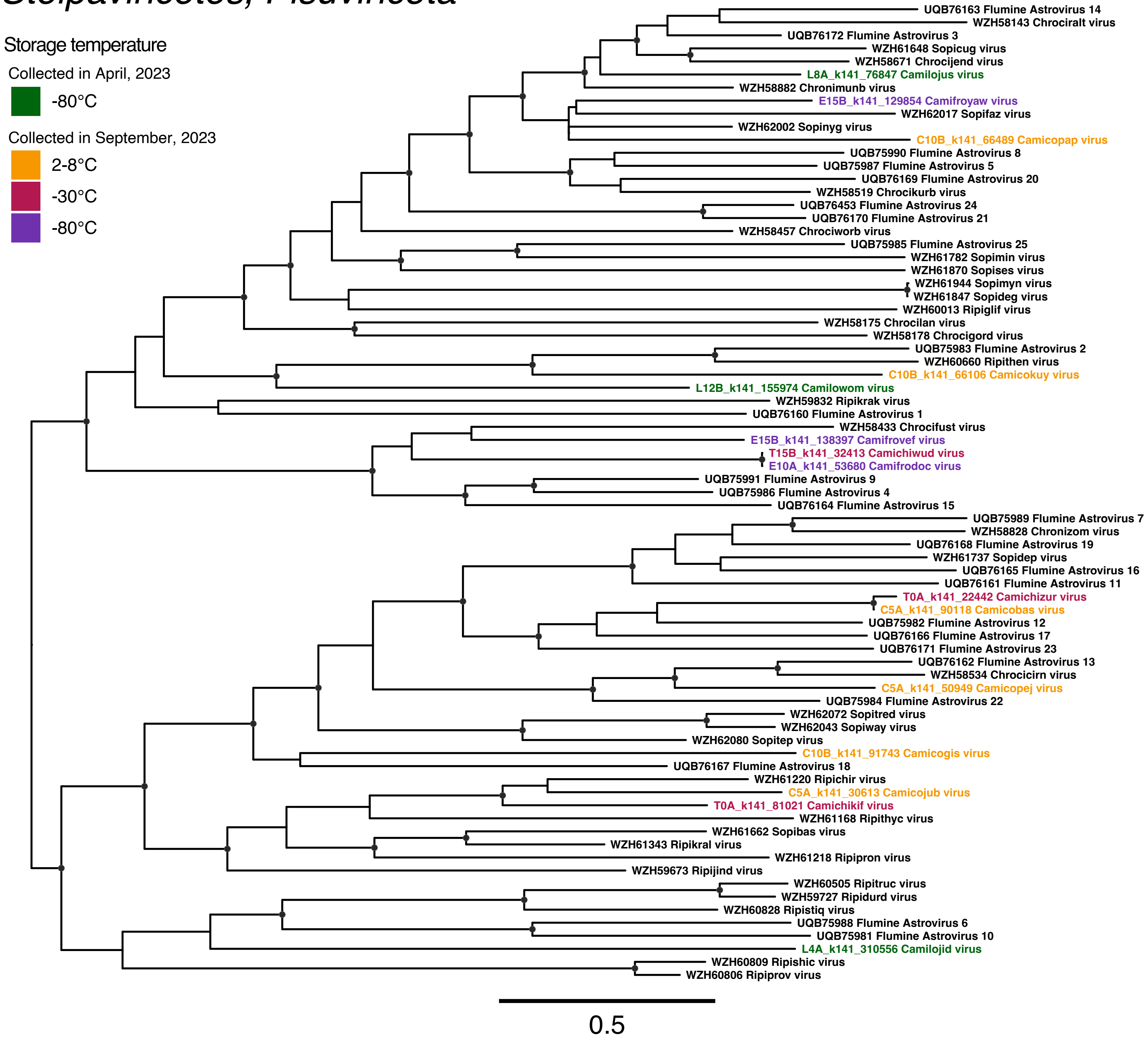

**Supplementary Figure 7.** Phylogenetic tree of RdRp amino acid sequences from the family *Astroviridae*. Putative novel viruses identified in this study are coloured by the storage temperature and collection month, such that samples collected in April 2023 (data set 1) and stored at -80°C are coloured in green, while samples collected in September 2023 (data set 2) and stored at 2-8°C are in orange, -30°C in red, and -80°C in purple. Known viruses are coloured in black. Trees are midpoint rooted with branch lengths scaled according to the number of amino acid substitutions per site. Black circles represent node support ≥80% using 1000 S H-aLRT replicates.

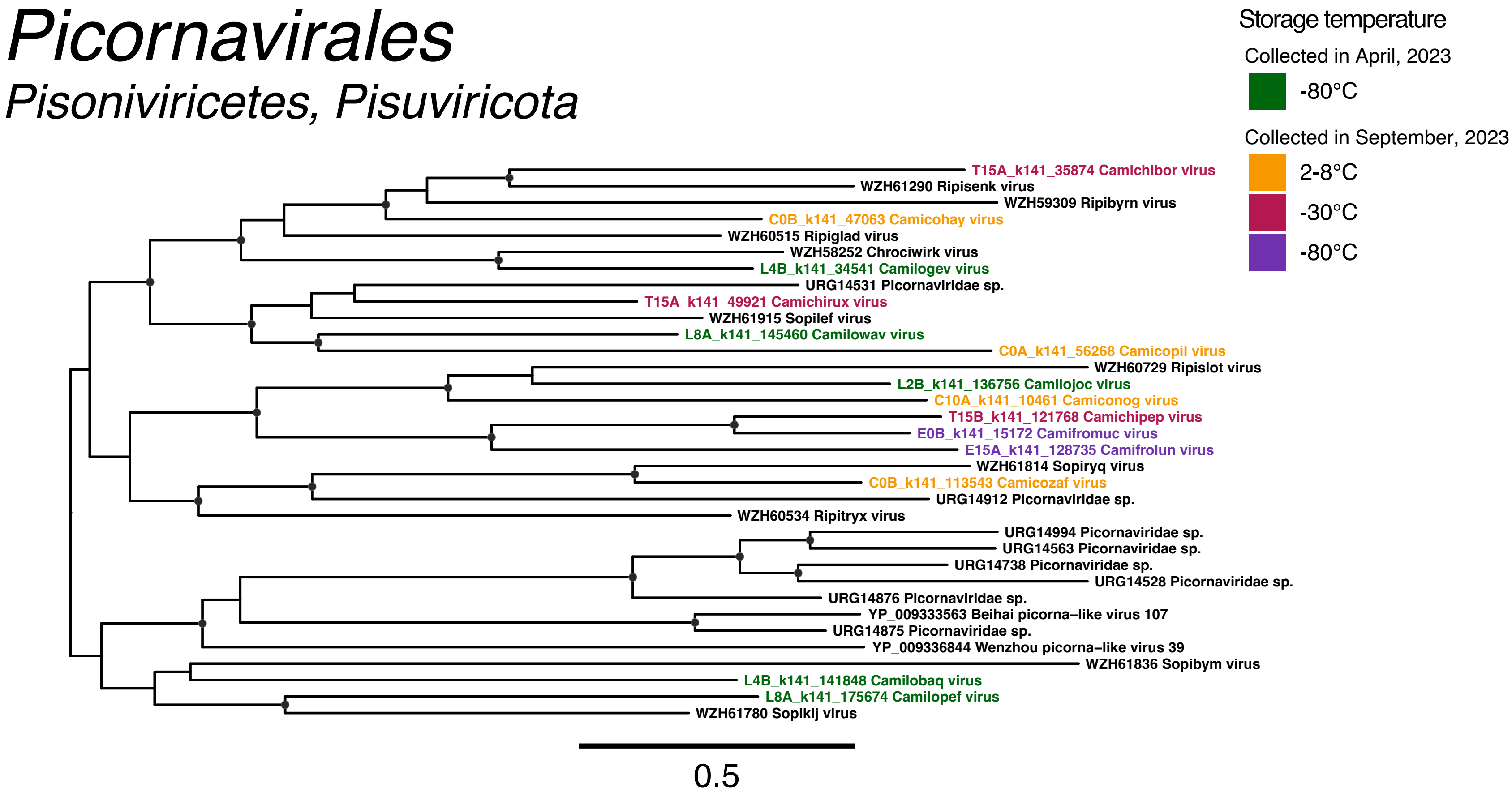

Picornavirales
Pisoniviricetes, Pisuviricota
Storage temperature
Collected in April, 2023
-80°C
Collected in September, 2023
2-8°C
-30°C
-80°C
T15A_k141_35874 Camichibor virus
WZH61290 Ripisenk virus
WZH59309 Ripibyrn virus
C0B_k141_47063 Camicohay virus
WZH60515 Ripiglad virus
WZH58252 Chrociwirk virus
L4B_k141_34541 Camilogev virus
URG14531 Picornaviridae sp.
T15A_k141_49921 Camichirux virus
WZH61915 Sopilef virus
L8A_k141_145460 Camilowav virus
C0A_k141_56268 Camicopil virus
WZH60729 Ripislot virus
L2B_k141_136756 Camilojoc virus
C10A_k141_10461 Camiconog virus
T15B_k141_121768 Camichipep virus
E0B_k141_15172 Camifromuc virus
E15A_k141_128735 Camifrolun virus
WZH61814 Sopiryq virus
C0B_k141_113543 Camicozaf virus
URG14912 Picornaviridae sp.
WZH60534 Ripitryx virus
URG14994 Picornaviridae sp.
URG14563 Picornaviridae sp.
URG14738 Picornaviridae sp.
URG14528 Picornaviridae sp.
URG14876 Picornaviridae sp.
YP_009333563 Beihai picorna–like virus 107
URG14875 Picornaviridae sp.
YP_009336844 Wenzhou picorna–like virus 39
WZH61836 Sopibym virus
L4B_k141_141848 Camilobaq virus
L8A_k141_175674 Camilopef virus
WZH61780 Sopikij virus
0.5

**Supplementary Figure 8.** Phylogenetic tree of RdRp amino acid sequences from the order *Picornavirales*. Putative novel viruses identified in this study are coloured by the storage temperature and collection month, such that samples collected in April 2023 (data set 1) and stored at -80°C are coloured in green, while samples collected in September 2023 (data set 2) and stored at 2-8°C are in orange, -30°C in red, and -80°C in purple. Known viruses are coloured in black. Trees are midpoint rooted with branch lengths scaled according to the number of amino acid substitutions per site. Black circles represent node support ≥80% using 1000 S H-aLRT replicates.

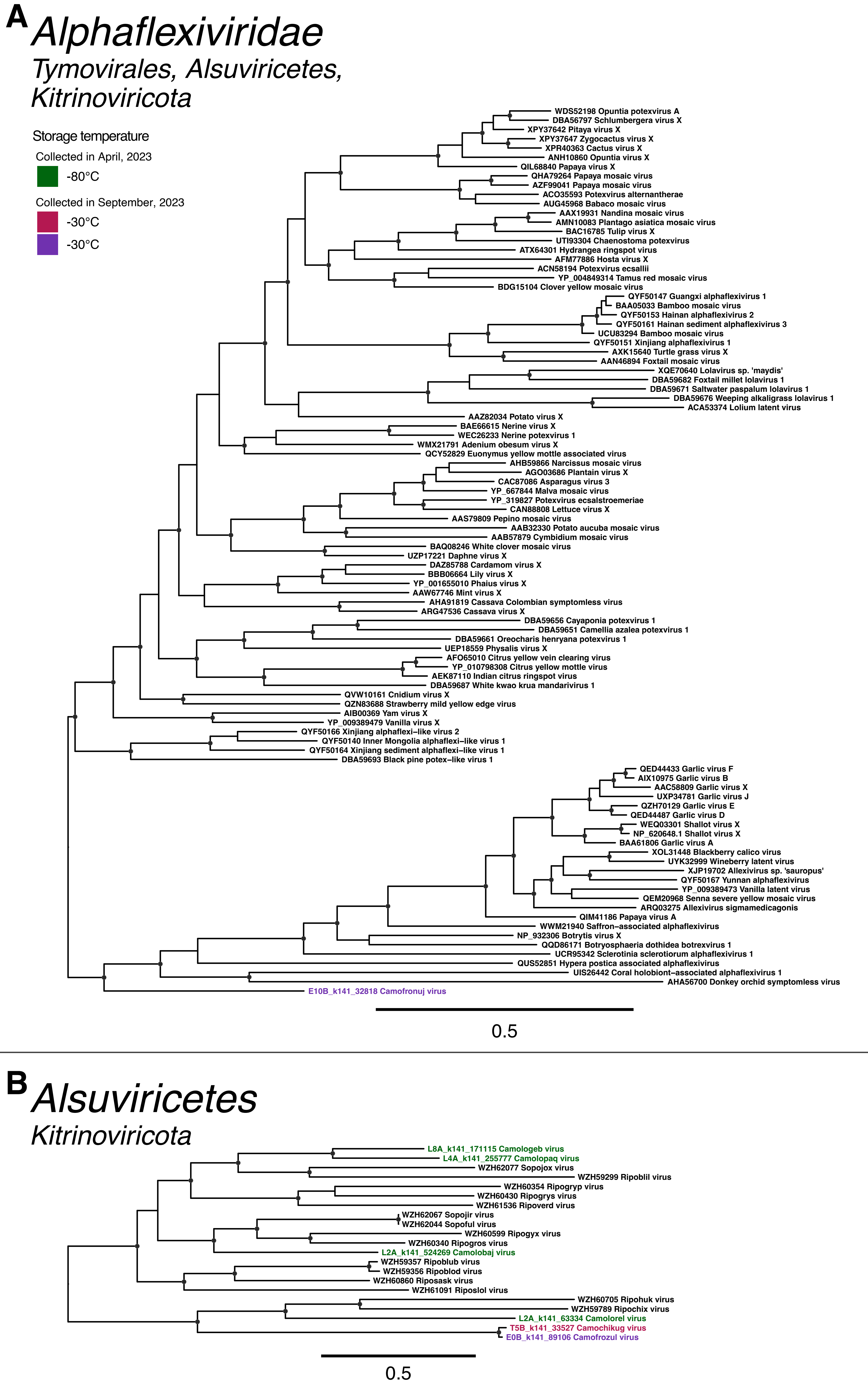
A
Alphaflexiviridae
Tymovirales, Alsuviricetes, Kitrinoviricota
Storage temperature
Collected in April, 2023
-80°C
Collected in September, 2023
-30°C
-30°C
WDS52198 Opuntia potexvirus A
DBA56797 Schlumbergera virus X
XPY37642 Pitaya virus X
XPY37647 Zygocactus virus X
XPR40363 Cactus virus X
ANH10860 Opuntia virus X
QIL68840 Papaya virus X
QHA79264 Papaya mosaic virus
AZF99041 Papaya mosaic virus
ACO35593 Potexvirus alternantherae
AUG45968 Babaco mosaic virus
AAX19931 Nandina mosaic virus
AMN10083 Plantago asiatica mosaic virus
BAC16785 Tulip virus X
UTI93304 Chaenostoma potexvirus
ATX64301 Hydrangea ringspot virus
AFM77886 Hosta virus X
ACN58194 Potexvirus ecsallii
YP_004849314 Tamus red mosaic virus
BDG15104 Clover yellow mosaic virus
QYF50147 Guangxi alphaflexivirus 1
BAA05033 Bamboo mosaic virus
QYF50153 Hainan alphaflexivirus 2
QYF50161 Hainan sediment alphaflexivirus 3
UCU83294 Bamboo mosaic virus
QYF50151 Xinjiang alphaflexivirus 1
AXK15640 Turtle grass virus X
AAN46894 Foxtail mosaic virus
XQE70640 Lolavirus sp. 'maydis'
DBA59682 Foxtail millet lolavirus 1
DBA59671 Saltwater paspalum lolavirus 1
DBA59676 Weeping alkaligrass lolavirus 1
ACA53374 Lolium latent virus
AAZ82034 Potato virus X
BAE66615 Nerine virus X
WEC26233 Nerine potexvirus 1
WMX21791 Adenium obesum virus X
QCY52829 Euonymus yellow mottle associated virus
AHB59866 Narcissus mosaic virus
AGO03686 Plantain virus X
CAC87086 Asparagus virus 3
YP_667844 Malva mosaic virus
YP_319827 Potexvirus ecsalstroemeriae
CAN88808 Lettuce virus X
AAS79809 Pepino mosaic virus
AAB32330 Potato aucuba mosaic virus
AAB57879 Cymbidium mosaic virus
BAQ08246 White clover mosaic virus
UZP17221 Daphne virus X
DAZ85788 Cardamom virus X
BBB06664 Lily virus X
YP_001655010 Phaius virus X
AAW67746 Mint virus X
AHA91819 Cassava Colombian symptomless virus
ARG47536 Cassava virus X
DBA59656 Cayaponia potexvirus 1
DBA59651 Camellia azalea potexvirus 1
DBA59661 Oreocharis henryana potexvirus 1
UEP18559 Physalis virus X
AFO65010 Citrus yellow vein clearing virus
YP_010798308 Citrus yellow mottle virus
AEK87110 Indian citrus ringspot virus
DBA59687 White kwao krua mandarivirus 1
QVW10161 Cnidium virus X
QZN83688 Strawberry mild yellow edge virus
AIB00369 Yam virus X
YP_009389479 Vanilla virus X
QYF50166 Xinjiang alphaflexi–like virus 2
QYF50140 Inner Mongolia alphaflexi–like virus 1
QYF50164 Xinjiang sediment alphaflexi–like virus 1
DBA59693 Black pine potex–like virus 1
QED44433 Garlic virus F
AIX10975 Garlic virus B
AAC58809 Garlic virus X
UXP34781 Garlic virus J
QZH70129 Garlic virus E
QED44487 Garlic virus D
WEQ03301 Shallot virus X
NP_620648.1 Shallot virus X
BAA61806 Garlic virus A
XOL31448 Blackberry calico virus
UYK32999 Wineberry latent virus
XJP19702 Allexivirus sp. 'sauropus'
QYF50167 Yunnan alphaflexivirus
YP_009389473 Vanilla latent virus
QEM20968 Senna severe yellow mosaic virus
ARQ03275 Allexivirus sigmamedicagonis
QIM41186 Papaya virus A
WWM21940 Saffron–associated alphaflexivirus
NP_932306 Botrytis virus X
QQD86171 Botryosphaeria dothidea botrexvirus 1
UCR95342 Sclerotinia sclerotiorum alphaflexivirus 1
QUS52851 Hypera postica associated alphaflexivirus
UIS26442 Coral holobiont–associated alphaflexivirus 1
AHA56700 Donkey orchid symptomless virus
E10B_k141_32818 Camofronuj virus
0.5
B
Alsuviricetes
Kitrinoviricota
L8A_k141_171115 Camologeb virus
L4A_k141_255777 Camolopaq virus
WZH62077 Sopojox virus
WZH59299 Ripoblil virus
WZH60354 Ripogryp virus
WZH60430 Ripogrys virus
WZH61536 Ripoverd virus
WZH62067 Sopojir virus
WZH62044 Sopoful virus
WZH60599 Ripogyx virus
WZH60340 Ripogros virus
L2A_k141_524269 Camolobaj virus
WZH59357 Ripoblub virus
WZH59356 Ripoblod virus
WZH60860 Riposask virus
WZH61091 Riposlol virus
WZH60705 Ripohuk virus
WZH59789 Ripochix virus
L2A_k141_63334 Camolorel virus
T5B_k141_33527 Camochikug virus
E0B_k141_89106 Camofrozul virus
0.5

**Supplementary Figure 9.** Phylogenetic tree of RdRp amino acid sequences from (A) the family *Alphaflexiviridae*, and (B) divergent *Alsuviricetes*-like sequences. Putative novel viruses identified in this study are coloured by the storage temperature and collection month, such that samples collected in April 2023 (data set 1) and stored at -80°C are coloured in green, while samples collected in September 2023 (data set 2) and stored at -30°C are in red and -80°C in purple. Known viruses are coloured in black. Trees are midpoint rooted with branch lengths scaled according to the number of amino acid substitutions per site. Black circles represent node support ≥80% using 1000 S H-aLRT replicates.

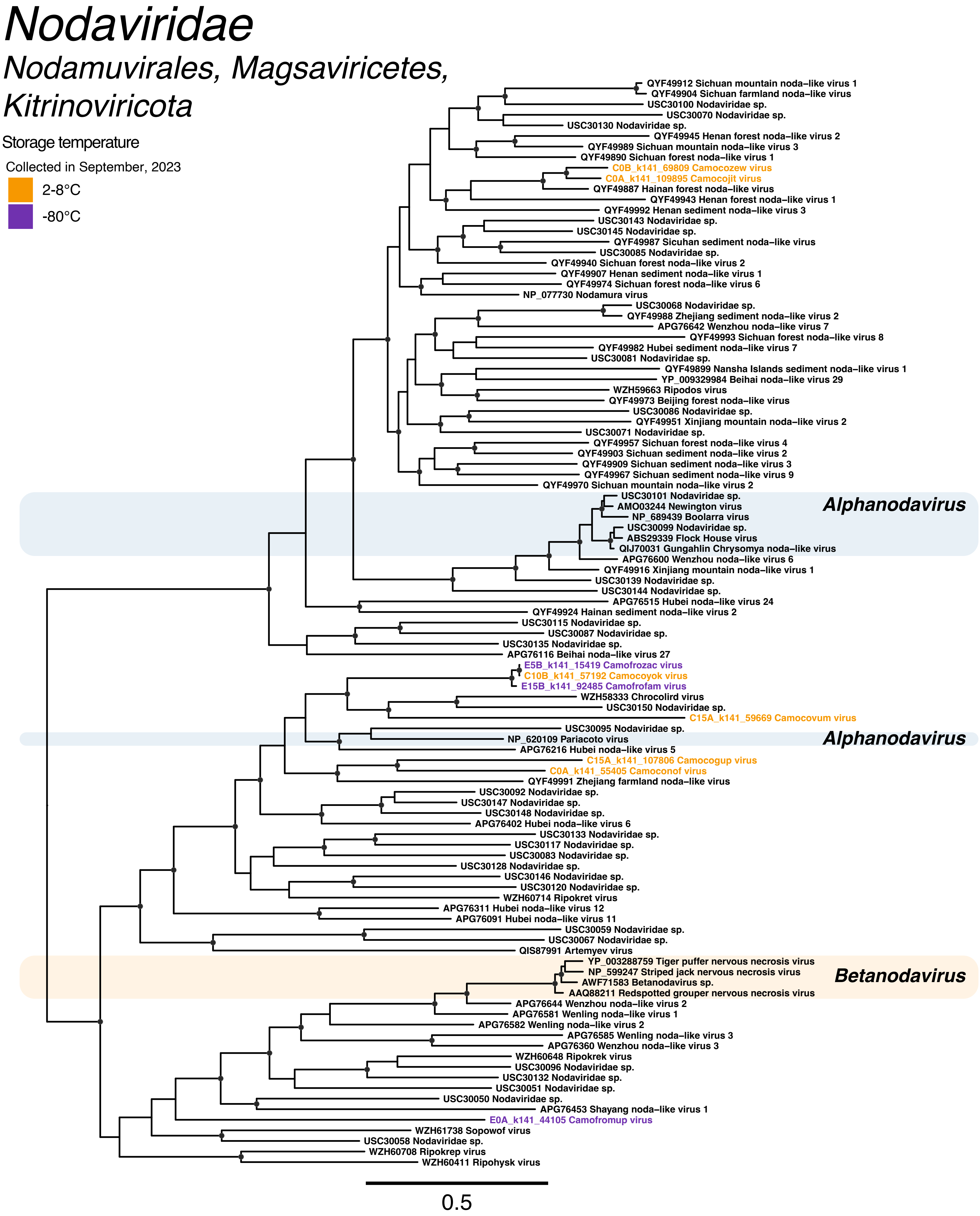

Nodaviridae
Nodamuvirales, Magsaviricetes, Kitrinoviricota
Storage temperature
Collected in September, 2023
2-8°C
-80°C
QYF49912 Sichuan mountain noda–like virus 1
QYF49904 Sichuan farmland noda–like virus
USC30100 Nodaviridae sp.
USC30070 Nodaviridae sp.
USC30130 Nodaviridae sp.
QYF49945 Henan forest noda–like virus 2
QYF49989 Sichuan mountain noda–like virus 3
QYF49890 Sichuan forest noda–like virus 1
C0B_k141_69809 Camocozew virus
C0A_k141_109895 Camocojit virus
QYF49887 Hainan forest noda–like virus
QYF49943 Henan forest noda–like virus 1
QYF49992 Henan sediment noda–like virus 3
USC30143 Nodaviridae sp.
USC30145 Nodaviridae sp.
QYF49987 Sicuhan sediment noda–like virus
USC30085 Nodaviridae sp.
QYF49940 Sichuan forest noda–like virus 2
QYF49907 Henan sediment noda–like virus 1
QYF49974 Sichuan forest noda–like virus 6
NP_077730 Nodamura virus
USC30068 Nodaviridae sp.
QYF49988 Zhejiang sediment noda–like virus 2
APG76642 Wenzhou noda–like virus 7
QYF49993 Sichuan forest noda–like virus 8
QYF49982 Hubei sediment noda–like virus 7
USC30081 Nodaviridae sp.
QYF49899 Nansha Islands sediment noda–like virus 1
YP_009329984 Beihai noda–like virus 29
WZH59663 Ripodos virus
QYF49973 Beijing forest noda–like virus
USC30086 Nodaviridae sp.
QYF49951 Xinjiang mountain noda–like virus 2
USC30071 Nodaviridae sp.
QYF49957 Sichuan forest noda–like virus 4
QYF49903 Sichuan sediment noda–like virus 2
QYF49909 Sichuan sediment noda–like virus 3
QYF49967 Sichuan sediment noda–like virus 9
QYF49970 Sichuan mountain noda–like virus 2
USC30101 Nodaviridae sp.
Alphanodavirus
AMO03244 Newington virus
NP_689439 Boolarra virus
USC30099 Nodaviridae sp.
ABS29339 Flock House virus
QIJ70031 Gungahlin Chrysomya noda–like virus
APG76600 Wenzhou noda–like virus 6
QYF49916 Xinjiang mountain noda–like virus 1
USC30139 Nodaviridae sp.
USC30144 Nodaviridae sp.
APG76515 Hubei noda–like virus 24
QYF49924 Hainan sediment noda–like virus 2
USC30115 Nodaviridae sp.
USC30087 Nodaviridae sp.
USC30135 Nodaviridae sp.
APG76116 Beihai noda–like virus 27
E5B_k141_15419 Camofrozac virus
C10B_k141_57192 Camocoyok virus
E15B_k141_92485 Camofrofam virus
WZH58333 Chrocolird virus
USC30150 Nodaviridae sp.
C15A_k141_59669 Camocovum virus
USC30095 Nodaviridae sp.
NP_620109 Pariacoto virus
Alphanodavirus
APG76216 Hubei noda–like virus 5
C15A_k141_107806 Camocogup virus
C0A_k141_55405 Camoconof virus
QYF49991 Zhejiang farmland noda–like virus
USC30092 Nodaviridae sp.
USC30147 Nodaviridae sp.
USC30148 Nodaviridae sp.
APG76402 Hubei noda–like virus 6
USC30133 Nodaviridae sp.
USC30117 Nodaviridae sp.
USC30083 Nodaviridae sp.
USC30128 Nodaviridae sp.
USC30146 Nodaviridae sp.
USC30120 Nodaviridae sp.
WZH60714 Ripokret virus
APG76311 Hubei noda–like virus 12
APG76091 Hubei noda–like virus 11
USC30059 Nodaviridae sp.
USC30067 Nodaviridae sp.
QIS87991 Artemyev virus
YP_003288759 Tiger puffer nervous necrosis virus
NP_599247 Striped jack nervous necrosis virus
Betanodavirus
AWF71583 Betanodavirus sp.
AAQ88211 Redspotted grouper nervous necrosis virus
APG76644 Wenzhou noda–like virus 2
APG76581 Wenling noda–like virus 1
APG76582 Wenling noda–like virus 2
APG76585 Wenling noda–like virus 3
APG76360 Wenzhou noda–like virus 3
WZH60648 Ripokrek virus
USC30096 Nodaviridae sp.
USC30132 Nodaviridae sp.
USC30051 Nodaviridae sp.
USC30050 Nodaviridae sp.
APG76453 Shayang noda–like virus 1
E0A_k141_44105 Camofromup virus
WZH61738 Sopowof virus
USC30058 Nodaviridae sp.
WZH60708 Ripokrep virus
WZH60411 Ripohysk virus
0.5

**Supplementary Figure 10.** Phylogenetic tree of RdRp amino acid sequences from the family *Nodaviridae*. Putative novel viruses identified in this study are coloured by the storage temperature and collection month, such that samples collected in September 2023 (data set 2) and stored at 2-8°C are coloured in orange and -80°C in purple. Known viruses are coloured in black. Trees are midpoint rooted with branch lengths scaled according to the number of amino acid substitutions per site. Black circles represent node support ≥80% using 1000 S H-aLRT replicates.

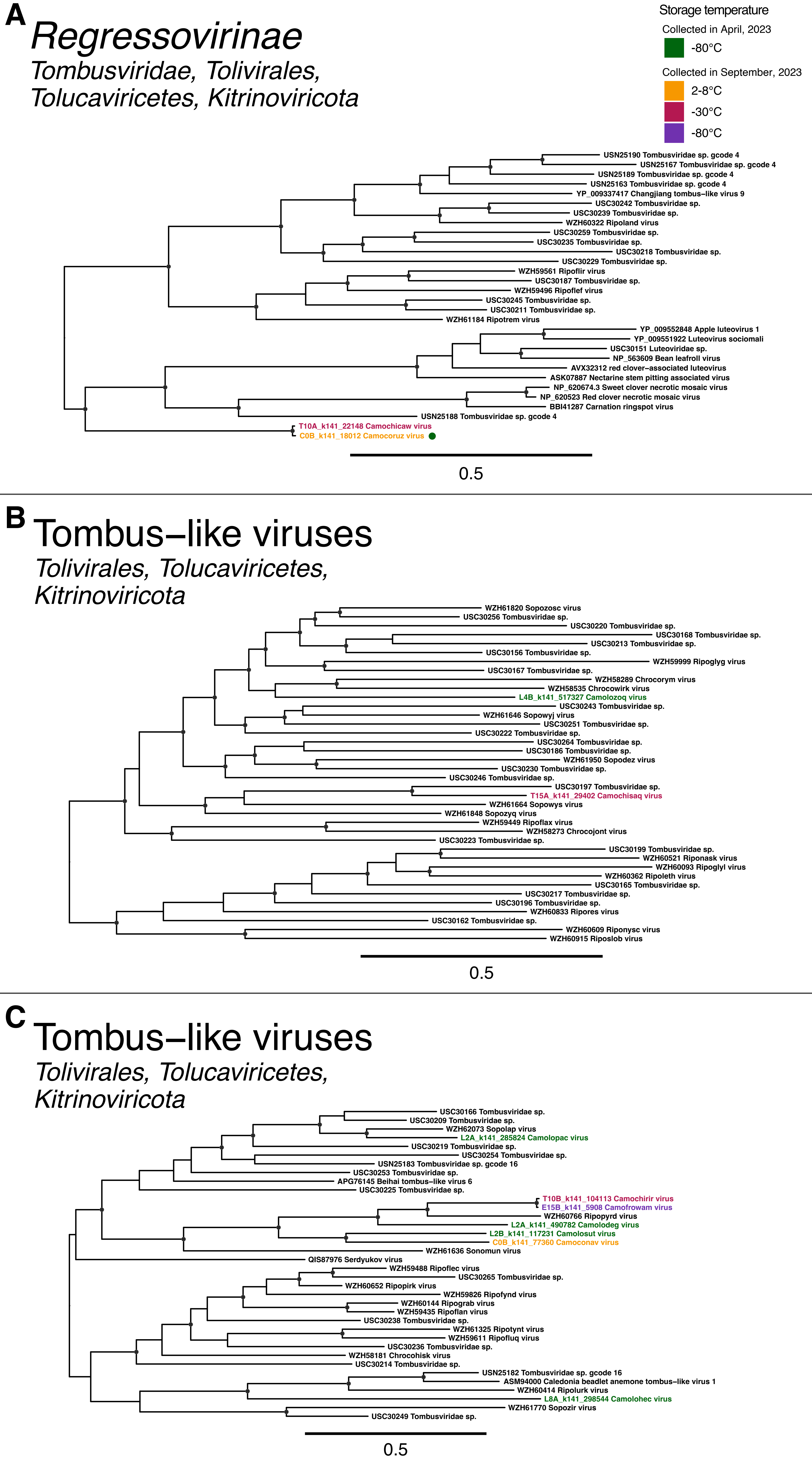
A
Regressovirinae
Tombusviridae, Tolivirales,
Tolucaviricetes, Kitrinoviricota
Storage temperature
Collected in April, 2023
-80°C
Collected in September, 2023
2-8°C
-30°C
-80°C
USN25190 Tombusviridae sp. gcode 4
USN25167 Tombusviridae sp. gcode 4
USN25189 Tombusviridae sp. gcode 4
USN25163 Tombusviridae sp. gcode 4
YP_009337417 Changjiang tombus-like virus 9
USC30242 Tombusviridae sp.
USC30239 Tombusviridae sp.
WZH60322 Ripoland virus
USC30259 Tombusviridae sp.
USC30235 Tombusviridae sp.
USC30218 Tombusviridae sp.
USC30229 Tombusviridae sp.
WZH59561 Ripoflir virus
USC30187 Tombusviridae sp.
WZH59496 Ripoflef virus
USC30245 Tombusviridae sp.
USC30211 Tombusviridae sp.
WZH61184 Ripotrem virus
YP_009552848 Apple luteovirus 1
YP_009551922 Luteovirus sociomali
USC30151 Luteoviridae sp.
NP_563609 Bean leafroll virus
AVX32312 red clover-associated luteovirus
ASK07887 Nectarine stem pitting associated virus
NP_620674.3 Sweet clover necrotic mosaic virus
NP_620523 Red clover necrotic mosaic virus
BBI41287 Carnation ringspot virus
USN25188 Tombusviridae sp. gcode 4
T10A_k141_22148 Camochicaw virus
C0B_k141_18012 Camocoruz virus
0.5
B
Tombus-like viruses
Tolivirales, Tolucaviricetes,
Kitrinoviricota
WZH61820 Sopozosc virus
USC30256 Tombusviridae sp.
USC30220 Tombusviridae sp.
USC30168 Tombusviridae sp.
USC30213 Tombusviridae sp.
USC30156 Tombusviridae sp.
WZH59999 Ripoglyg virus
USC30167 Tombusviridae sp.
WZH58289 Chrocorym virus
WZH58535 Chrocowirk virus
L4B_k141_517327 Camolozoq virus
USC30243 Tombusviridae sp.
WZH61646 Sopowyj virus
USC30251 Tombusviridae sp.
USC30222 Tombusviridae sp.
USC30264 Tombusviridae sp.
USC30186 Tombusviridae sp.
WZH61950 Sopodez virus
USC30230 Tombusviridae sp.
USC30246 Tombusviridae sp.
USC30197 Tombusviridae sp.
T15A_k141_29402 Camochisaq virus
WZH61664 Sopowys virus
WZH61848 Sopozyq virus
WZH59449 Ripoflax virus
WZH58273 Chrocojont virus
USC30223 Tombusviridae sp.
USC30199 Tombusviridae sp.
WZH60521 Riponask virus
WZH60093 Ripoglyl virus
WZH60362 Ripoleth virus
USC30165 Tombusviridae sp.
USC30217 Tombusviridae sp.
USC30196 Tombusviridae sp.
WZH60833 Ripores virus
USC30162 Tombusviridae sp.
WZH60609 Riponysc virus
WZH60915 Riposlob virus
0.5
C
Tombus-like viruses
Tolivirales, Tolucaviricetes,
Kitrinoviricota
USC30166 Tombusviridae sp.
USC30209 Tombusviridae sp.
WZH62073 Sopolap virus
L2A_k141_285824 Camolopac virus
USC30219 Tombusviridae sp.
USC30254 Tombusviridae sp.
USN25183 Tombusviridae sp. gcode 16
USC30253 Tombusviridae sp.
APG76145 Beihai tombus-like virus 6
USC30225 Tombusviridae sp.
T10B_k141_104113 Camochirir virus
E15B_k141_5908 Camofrowam virus
WZH60766 Ripopyrd virus
L2A_k141_490782 Camolodeg virus
L2B_k141_117231 Camolosut virus
C0B_k141_77360 Camoconav virus
WZH61636 Sonomun virus
QIS87976 Serdyukov virus
WZH59488 Ripoflec virus
USC30265 Tombusviridae sp.
WZH60652 Ripopirk virus
WZH59826 Ripofynd virus
WZH60144 Ripograb virus
WZH59435 Ripoflan virus
USC30238 Tombusviridae sp.
WZH61325 Ripotynt virus
WZH59611 Ripofluq virus
USC30236 Tombusviridae sp.
WZH58181 Chrocohisk virus
USC30214 Tombusviridae sp.
USN25182 Tombusviridae sp. gcode 16
ASM94000 Caledonia beadlet anemone tombus-like virus 1
WZH60414 Ripolurk virus
L8A_k141_298544 Camolohec virus
WZH61770 Sopozir virus
USC30249 Tombusviridae sp.
0.5

**Supplementary Figure 11.** Phylogenetic tree of RdRp amino acid sequences from (A) the subfamily *Regressovirinae*, and (B, C) two distinct groups of divergent tombus-like viruses. Putative novel viruses identified in this study are coloured by the storage temperature and collection month, such that samples collected in April 2023 (data set 1) and stored at -80°C are coloured in green, while samples collected in September 2023 (data set 2) and stored at 2-8°C are in orange, -30°C in red, and -80°C in purple. Known viruses are coloured in black. Sequences found in both data sets are indicated by a coloured circle next to the provisional virus name, coloured by the storage temperature of library or libraries the identical sequence was found in. Trees are midpoint rooted with branch lengths scaled according to the number of amino acid substitutions per site. Black circles represent node support ≥80% using 1000 S H-aLRT replicates.

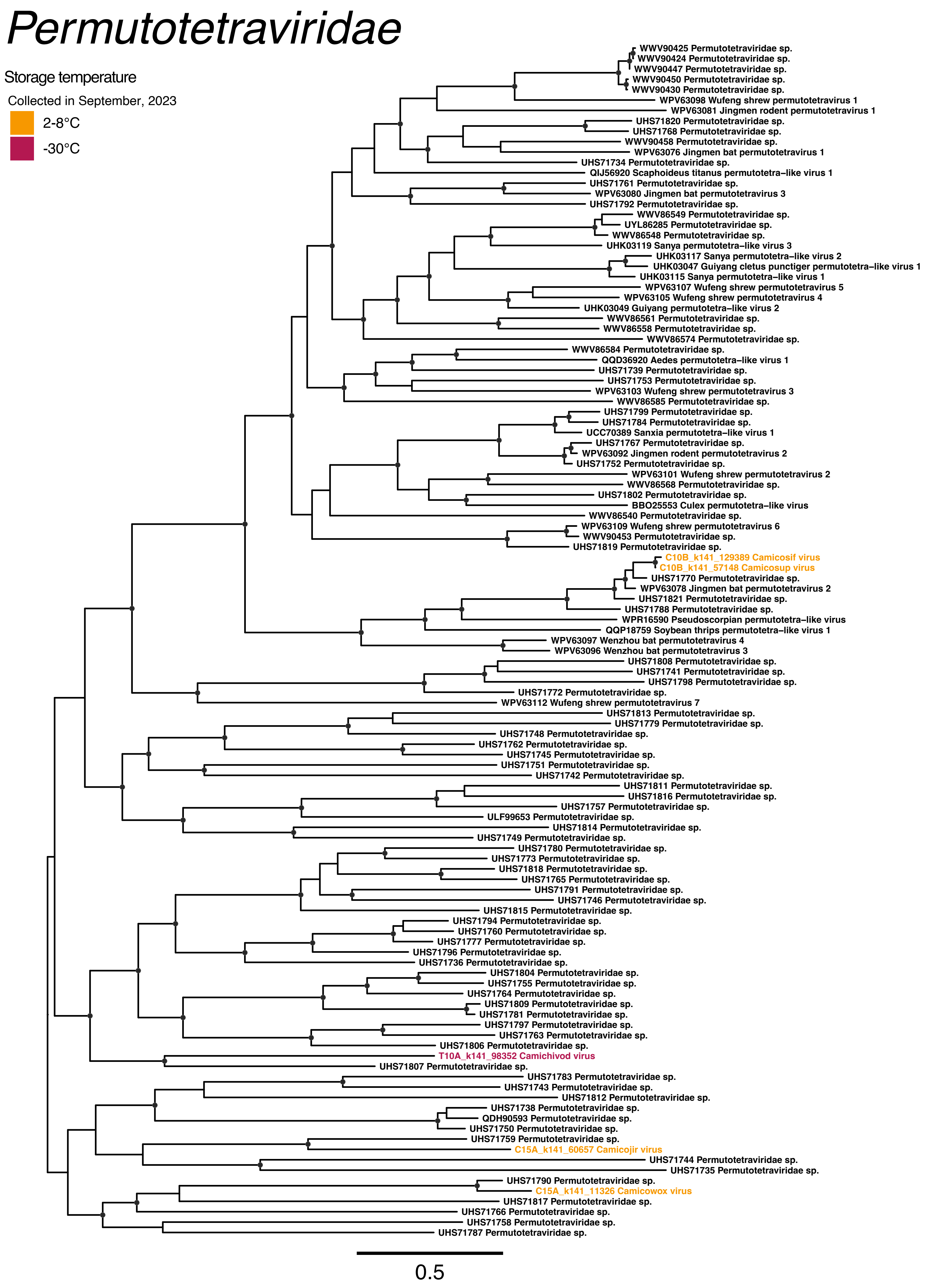
Permutotetraviridae
Storage temperature
Collected in September, 2023
2-8°C
-30°C
WWV90425 Permutotetraviridae sp.
WWV90424 Permutotetraviridae sp.
WWV90447 Permutotetraviridae sp.
WWV90450 Permutotetraviridae sp.
WWV90430 Permutotetraviridae sp.
WPV63098 Wufeng shrew permutotetravirus 1
WPV63081 Jingmen rodent permutotetravirus 1
UHS71820 Permutotetraviridae sp.
UHS71768 Permutotetraviridae sp.
WWV90458 Permutotetraviridae sp.
WPV63076 Jingmen bat permutotetravirus 1
UHS71734 Permutotetraviridae sp.
QIJ56920 Scaphoideus titanus permutotetra−like virus 1
UHS71761 Permutotetraviridae sp.
WPV63080 Jingmen bat permutotetravirus 3
UHS71792 Permutotetraviridae sp.
WWV86549 Permutotetraviridae sp.
UYL86285 Permutotetraviridae sp.
WWV86548 Permutotetraviridae sp.
UHK03119 Sanya permutotetra−like virus 3
UHK03117 Sanya permutotetra−like virus 2
UHK03047 Guiyang cletus punctiger permutotetra−like virus 1
UHK03115 Sanya permutotetra−like virus 1
WPV63107 Wufeng shrew permutotetravirus 5
WPV63105 Wufeng shrew permutotetravirus 4
UHK03049 Guiyang permutotetra−like virus 2
WWV86561 Permutotetraviridae sp.
WWV86558 Permutotetraviridae sp.
WWV86574 Permutotetraviridae sp.
WWV86584 Permutotetraviridae sp.
QQD36920 Aedes permutotetra−like virus 1
UHS71739 Permutotetraviridae sp.
UHS71753 Permutotetraviridae sp.
WPV63103 Wufeng shrew permutotetravirus 3
WWV86585 Permutotetraviridae sp.
UHS71799 Permutotetraviridae sp.
UHS71784 Permutotetraviridae sp.
UCC70389 Sanxia permutotetra−like virus 1
UHS71767 Permutotetraviridae sp.
WPV63092 Jingmen rodent permutotetravirus 2
UHS71752 Permutotetraviridae sp.
WPV63101 Wufeng shrew permutotetravirus 2
WWV86568 Permutotetraviridae sp.
UHS71802 Permutotetraviridae sp.
BBO25553 Culex permutotetra−like virus
WWV86540 Permutotetraviridae sp.
WPV63109 Wufeng shrew permutotetravirus 6
WWV90453 Permutotetraviridae sp.
UHS71819 Permutotetraviridae sp.
C10B_k141_129389 Camicosif virus
C10B_k141_57148 Camicosup virus
UHS71770 Permutotetraviridae sp.
WPV63078 Jingmen bat permutotetravirus 2
UHS71821 Permutotetraviridae sp.
UHS71788 Permutotetraviridae sp.
WPR16590 Pseudoscorpian permutotetra−like virus
QQP18759 Soybean thrips permutotetra−like virus 1
WPV63097 Wenzhou bat permutotetravirus 4
WPV63096 Wenzhou bat permutotetravirus 3
UHS71808 Permutotetraviridae sp.
UHS71741 Permutotetraviridae sp.
UHS71798 Permutotetraviridae sp.
UHS71772 Permutotetraviridae sp.
WPV63112 Wufeng shrew permutotetravirus 7
UHS71813 Permutotetraviridae sp.
UHS71779 Permutotetraviridae sp.
UHS71748 Permutotetraviridae sp.
UHS71762 Permutotetraviridae sp.
UHS71745 Permutotetraviridae sp.
UHS71751 Permutotetraviridae sp.
UHS71742 Permutotetraviridae sp.
UHS71811 Permutotetraviridae sp.
UHS71816 Permutotetraviridae sp.
UHS71757 Permutotetraviridae sp.
ULF99653 Permutotetraviridae sp.
UHS71814 Permutotetraviridae sp.
UHS71749 Permutotetraviridae sp.
UHS71780 Permutotetraviridae sp.
UHS71773 Permutotetraviridae sp.
UHS71818 Permutotetraviridae sp.
UHS71765 Permutotetraviridae sp.
UHS71791 Permutotetraviridae sp.
UHS71746 Permutotetraviridae sp.
UHS71815 Permutotetraviridae sp.
UHS71794 Permutotetraviridae sp.
UHS71760 Permutotetraviridae sp.
UHS71777 Permutotetraviridae sp.
UHS71796 Permutotetraviridae sp.
UHS71736 Permutotetraviridae sp.
UHS71804 Permutotetraviridae sp.
UHS71755 Permutotetraviridae sp.
UHS71764 Permutotetraviridae sp.
UHS71809 Permutotetraviridae sp.
UHS71781 Permutotetraviridae sp.
UHS71797 Permutotetraviridae sp.
UHS71763 Permutotetraviridae sp.
UHS71806 Permutotetraviridae sp.
T10A_k141_98352 Camichivod virus
UHS71807 Permutotetraviridae sp.
UHS71783 Permutotetraviridae sp.
UHS71743 Permutotetraviridae sp.
UHS71812 Permutotetraviridae sp.
UHS71738 Permutotetraviridae sp.
QDH90593 Permutotetraviridae sp.
UHS71750 Permutotetraviridae sp.
UHS71759 Permutotetraviridae sp.
C15A_k141_60657 Camicojir virus
UHS71744 Permutotetraviridae sp.
UHS71735 Permutotetraviridae sp.
UHS71790 Permutotetraviridae sp.
C15A_k141_11326 Camicowox virus
UHS71817 Permutotetraviridae sp.
UHS71766 Permutotetraviridae sp.
UHS71758 Permutotetraviridae sp.
UHS71787 Permutotetraviridae sp.
0.5

**Supplementary Figure 12.** Phylogenetic tree of RdRp amino acid sequences from the family *Permutotetraviridae*. Putative novel viruses identified in this study are coloured by the storage temperature and collection month, such that samples collected in September 2023 (data set 2) and stored at 2-8°C are coloured in orange and -30°C in red. Known viruses are coloured in black. Trees are midpoint rooted with branch lengths scaled according to the number of amino acid substitutions per site. Black circles represent node support ≥80% using 1000 S H-aLRT replicates.

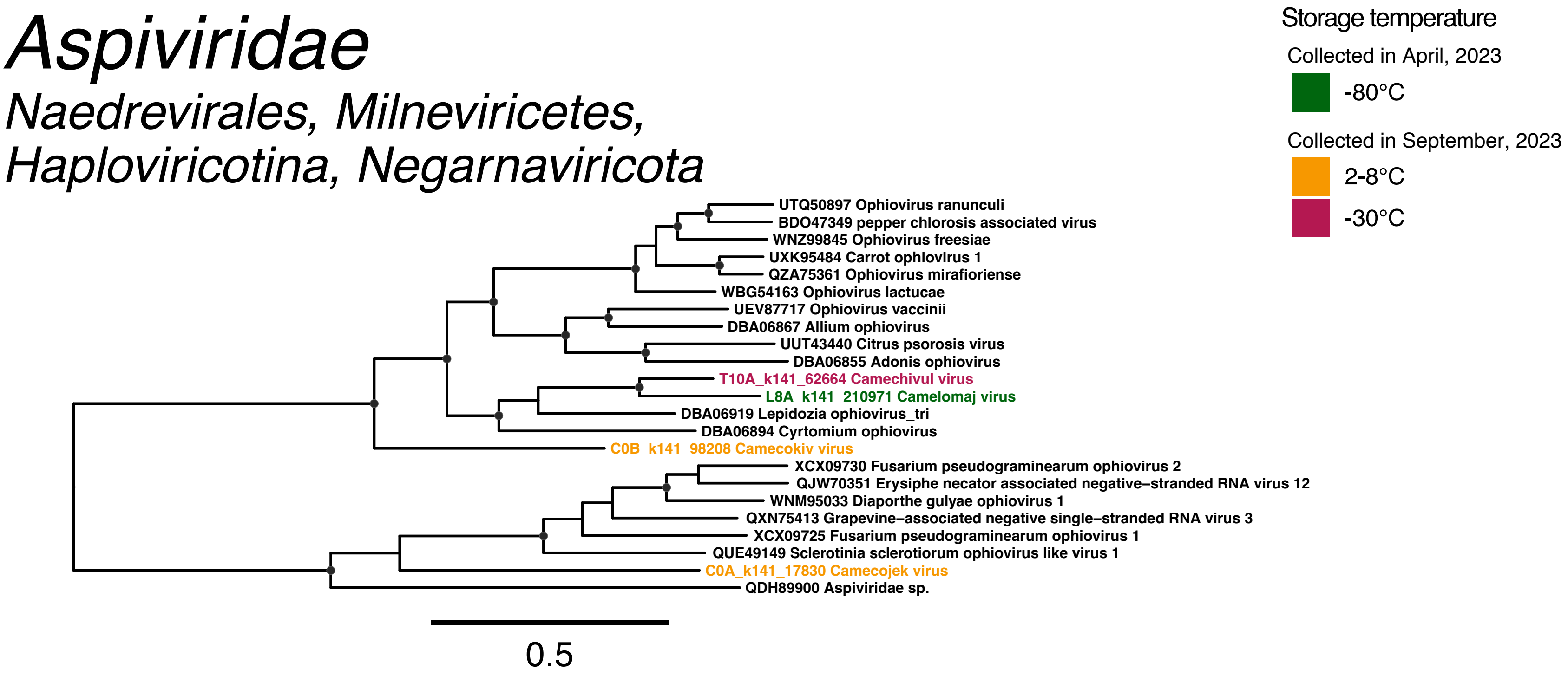
Aspiviridae
Naedrevirales, Milneviricetes,
Haploviricotina, Negarnaviricota
Storage temperature
Collected in April, 2023
-80°C
Collected in September, 2023
2-8°C
-30°C
UTQ50897 Ophiovirus ranunculi
BDO47349 pepper chlorosis associated virus
WNZ99845 Ophiovirus freesiae
UXK95484 Carrot ophiovirus 1
QZA75361 Ophiovirus mirafioriense
WBG54163 Ophiovirus lactucae
UEV87717 Ophiovirus vaccinii
DBA06867 Allium ophiovirus
UUT43440 Citrus psorosis virus
DBA06855 Adonis ophiovirus
T10A_k141_62664 Camechivul virus
L8A_k141_210971 Camelomaj virus
DBA06919 Lepidozia ophiovirus_tri
DBA06894 Cyrtomium ophiovirus
C0B_k141_98208 Camecokiv virus
XCX09730 Fusarium pseudograminearum ophiovirus 2
QJW70351 Erysiphe necator associated negative–stranded RNA virus 12
WNM95033 Diaporthe gulyae ophiovirus 1
QXN75413 Grapevine–associated negative single–stranded RNA virus 3
XCX09725 Fusarium pseudograminearum ophiovirus 1
QUE49149 Sclerotinia sclerotiorum ophiovirus like virus 1
C0A_k141_17830 Camecojek virus
QDH89900 Aspiviridae sp.
0.5

**Supplementary Figure 13.** Phylogenetic tree of RdRp amino acid sequences from the family *Aspiviridae*. Putative novel viruses identified in this study are coloured by the storage temperature and collection month, such that samples collected in April 2023 (data set 1) and stored at -80°C are coloured in green, while samples collected in September 2023 (data set 2) and stored at 2-8°C are in orange and -30°C in red. Known viruses are coloured in black. Trees are midpoint rooted with branch lengths scaled according to the number of amino acid substitutions per site. Black circles represent node support ≥80% using 1000 S H-aLRT replicates.

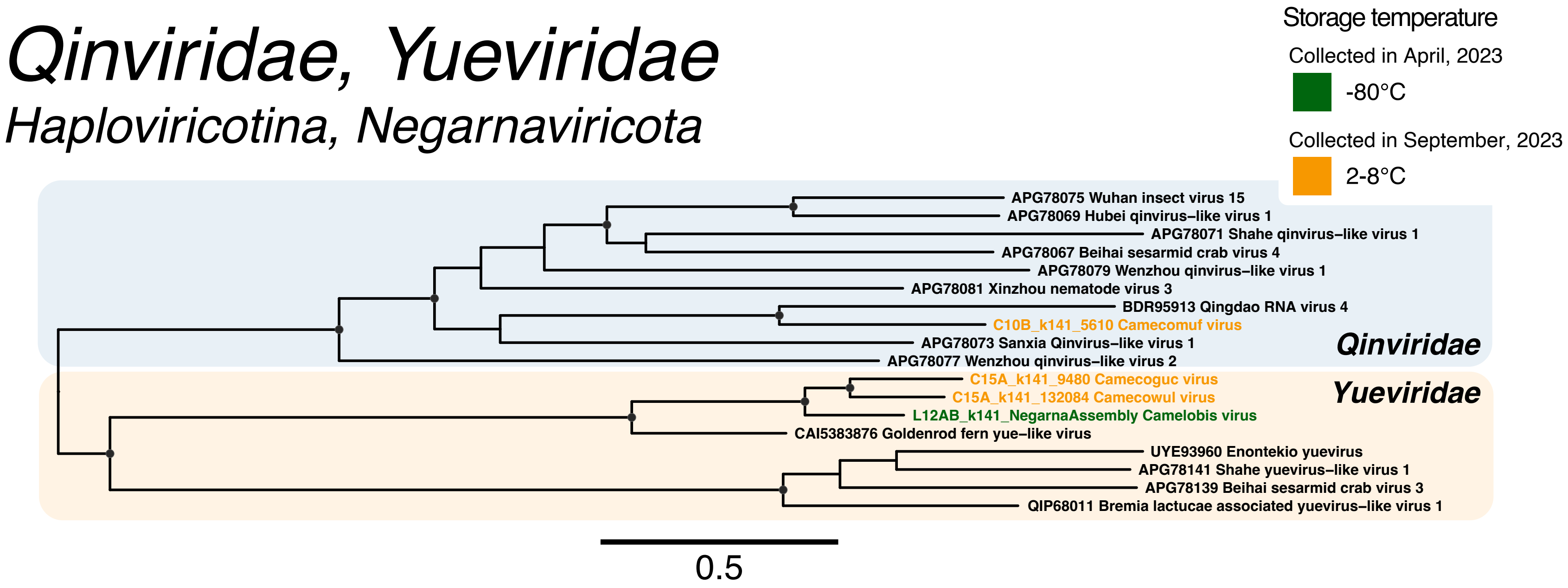
Qinviridae, Yueviridae
Haploviricotina, Negarnaviricota
Storage temperature
Collected in April, 2023
-80°C
Collected in September, 2023
2-8°C
APG78075 Wuhan insect virus 15
APG78069 Hubei qinvirus–like virus 1
APG78071 Shahe qinvirus–like virus 1
APG78067 Beihai sesarmid crab virus 4
APG78079 Wenzhou qinvirus–like virus 1
APG78081 Xinzhou nematode virus 3
BDR95913 Qingdao RNA virus 4
C10B_k141_5610 Camecomuf virus
APG78073 Sanxia Qinvirus–like virus 1
APG78077 Wenzhou qinvirus–like virus 2
Qinviridae
C15A_k141_9480 Camecoguc virus
C15A_k141_132084 Camecowul virus
L12AB_k141_NegarnaAssembly Camelobis virus
CAI5383876 Goldenrod fern yue–like virus
Yueviridae
UYE93960 Enontekio yuevirus
APG78141 Shahe yuevirus–like virus 1
APG78139 Beihai sesarmid crab virus 3
QIP68011 Bremia lactucae associated yuevirus–like virus 1
0.5

**Supplementary Figure 14.** Phylogenetic tree of RdRp amino acid sequences from the families *Qinviridae* and *Yueviridae*. Putative novel viruses identified in this study are coloured by the storage temperature and collection month, such that samples collected in April 2023 (data set 1) and stored at -80°C are coloured in green, while samples collected in September 2023 (data set 2) and stored at 2-8°C are in orange. Known viruses are coloured in black. Trees are midpoint rooted with branch lengths scaled according to the number of amino acid substitutions per site. Black circles represent node support ≥80% using 1000 S H-aLRT replicates.

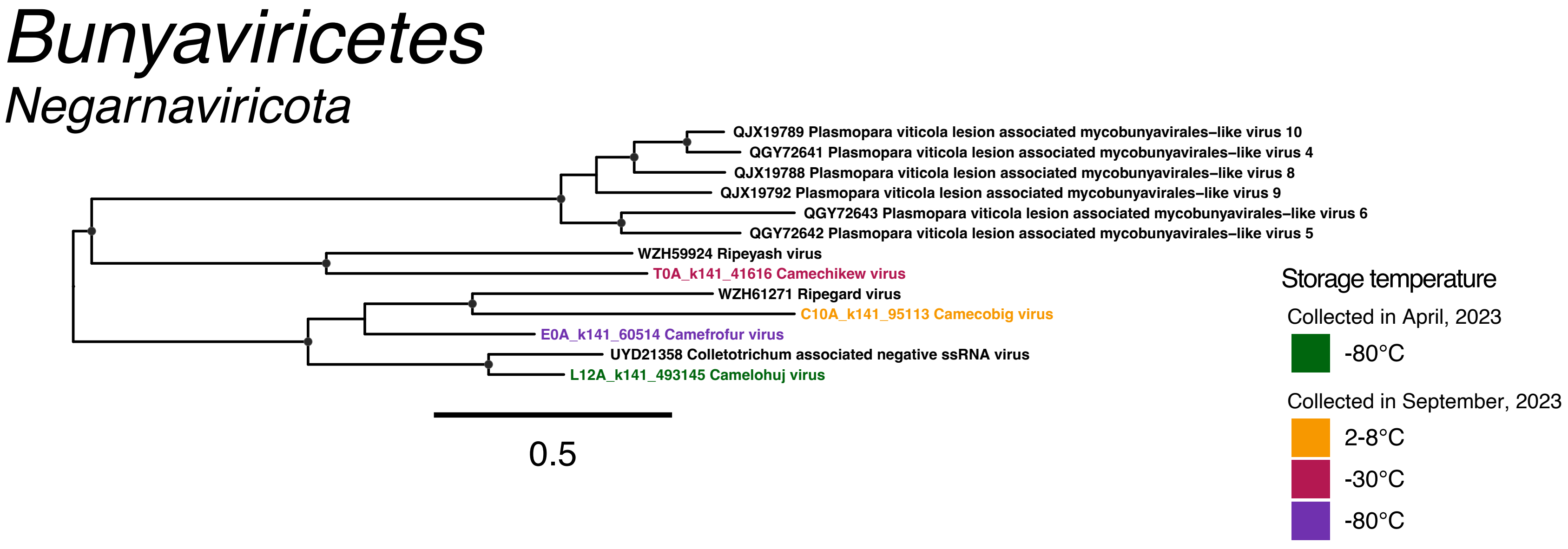
Bunyaviricetes
Negarnaviricota
QJX19789 Plasmopara viticola lesion associated mycobunyavirales–like virus 10
QGY72641 Plasmopara viticola lesion associated mycobunyavirales–like virus 4
QJX19788 Plasmopara viticola lesion associated mycobunyavirales–like virus 8
QJX19792 Plasmopara viticola lesion associated mycobunyavirales–like virus 9
QGY72643 Plasmopara viticola lesion associated mycobunyavirales–like virus 6
QGY72642 Plasmopara viticola lesion associated mycobunyavirales–like virus 5
WZH59924 Ripeyash virus
T0A_k141_41616 Camechikew virus
WZH61271 Ripegard virus
C10A_k141_95113 Camecobig virus
E0A_k141_60514 Camefrofur virus
UYD21358 Colletotrichum associated negative ssRNA virus
L12A_k141_493145 Camelohuj virus
0.5
Storage temperature
Collected in April, 2023
-80°C
Collected in September, 2023
2-8°C
-30°C
-80°C

**Supplementary Figure 15.** Phylogenetic tree of RdRp amino acid sequences from the class *Bunyaviricetes*. Putative novel viruses identified in this study are coloured by the storage temperature and collection month, such that samples collected in April 2023 (data set 1) and stored at -80°C are coloured in green, while samples collected in September 2023 (data set 2) and stored at 2-8°C are in orange, -30°C in red, and -80°C in purple. Known viruses are coloured in black. Trees are midpoint rooted with branch lengths scaled according to the number of amino acid substitutions per site. Black circles represent node support ≥80% using 1000 S H-aLRT replicates.

### *Amalgaviridae*

#### *Durnavirales, Duplopiviricetes, Pisuviricota*

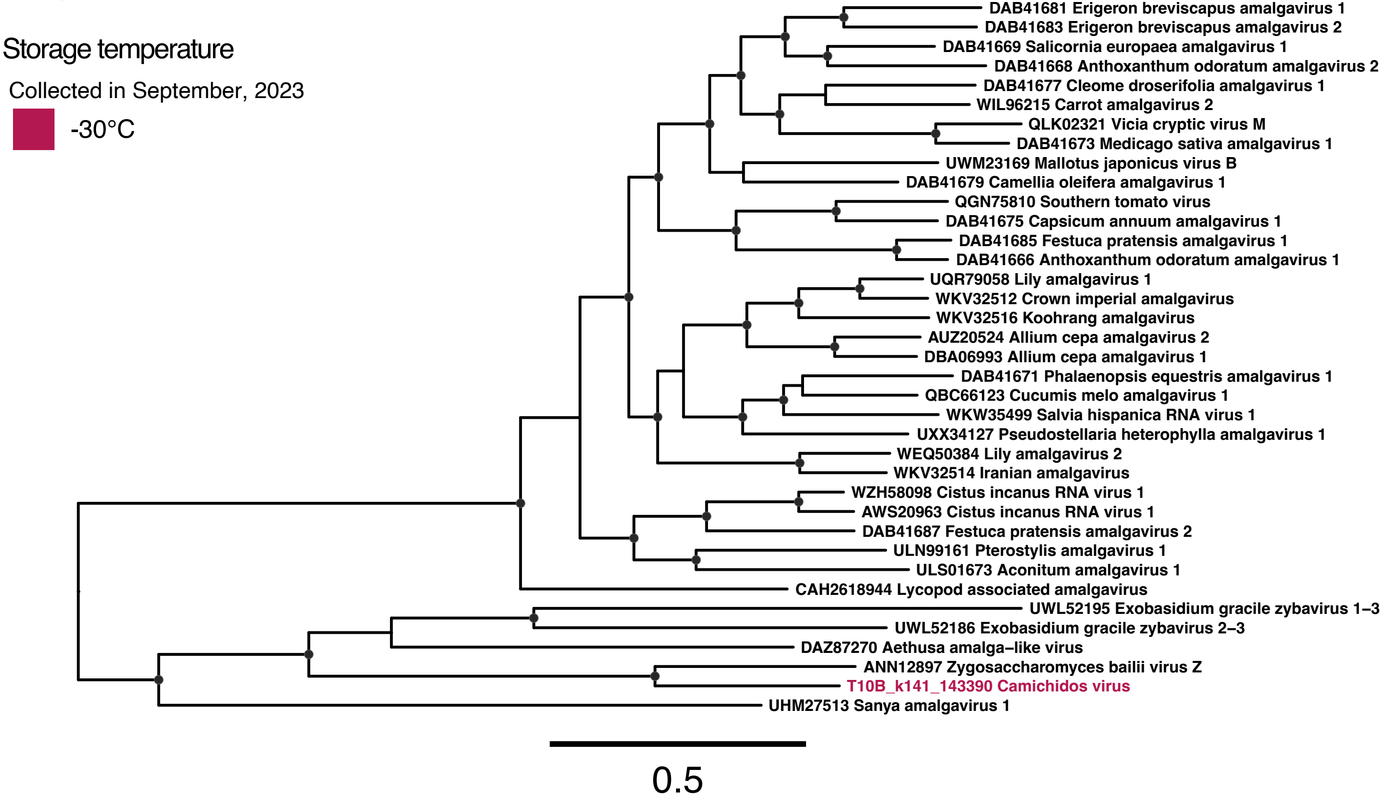

**Supplementary Figure 16.** Phylogenetic tree of RdRp amino acid sequences from the family *Amalgaviridae*. Putative novel viruses identified in this study are coloured by the storage temperature and collection month, such that samples collected in September 2023 (data set 2) and stored at -30°C are coloured in red. Known viruses are coloured in black. Trees are midpoint rooted with branch lengths scaled according to the number of amino acid substitutions per site. Black circles represent node support ≥80% using 1000 S H-aLRT replicates.

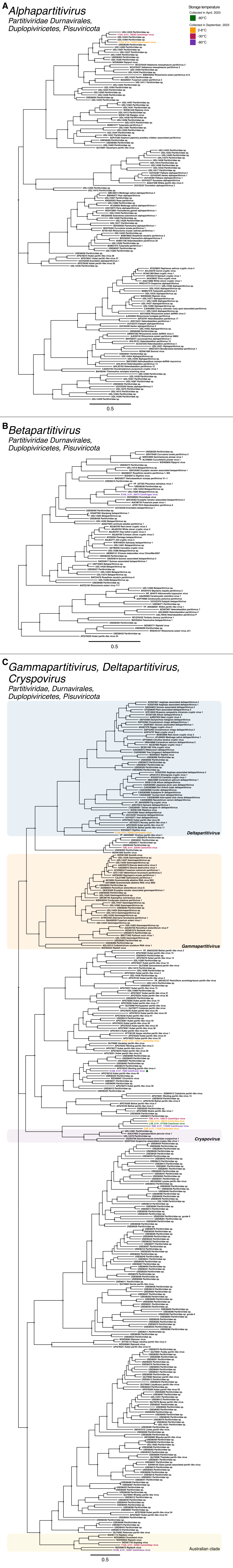
A
Alphapartitivirus
Partitiviridae Durnavirales, Duplopiviricetes, Pisuviricota
Storage temperature
Collected in April, 2023
-80°C
Collected in September, 2023
2-8°C
-30°C
-80°C
0.5
B
Betapartitivirus
Partitiviridae Durnavirales, Duplopiviricetes, Pisuviricota
0.5
C
Gammapartitivirus, Deltapartitivirus, Cryspovirus
Partitiviridae, Durnavirales, Duplopiviricetes, Pisuviricota
Deltapartitivirus
Gammapartitivirus
Cryspovirus
Australian clade
0.5

**Supplementary Figure 17.** Phylogenetic tree of RdRp amino acid sequences from (A) the genus *Alphapartitivirus*, (B) the genus *Betapartitiviridae*, and (C) the genera *Gammapartitivirus, Deltapartitiviridae*, and *Cryspovirus*, and divergent partiti-like sequences. Putative novel viruses identified in this study are coloured by the storage temperature and collection month, such that samples collected in April 2023 (data set 1) and stored at -80°C are coloured in green, while samples collected in September 2023 (data set 2) and stored at 2-8°C are in orange, -30°C in red, and -80°C in purple. Known viruses are coloured in black. Sequences found in both data sets are indicated by a coloured circle next to the provisional virus name, coloured by the storage temperature of library or libraries the identical sequence was found in. Clades comprised exclusively of sequences sourced from Australian environmental samples are indicated by a yellow box. Trees are midpoint rooted with branch lengths scaled according to the number of amino acid substitutions per site. Black circles represent node support ≥80% using 1000 S H-aLRT replicates.

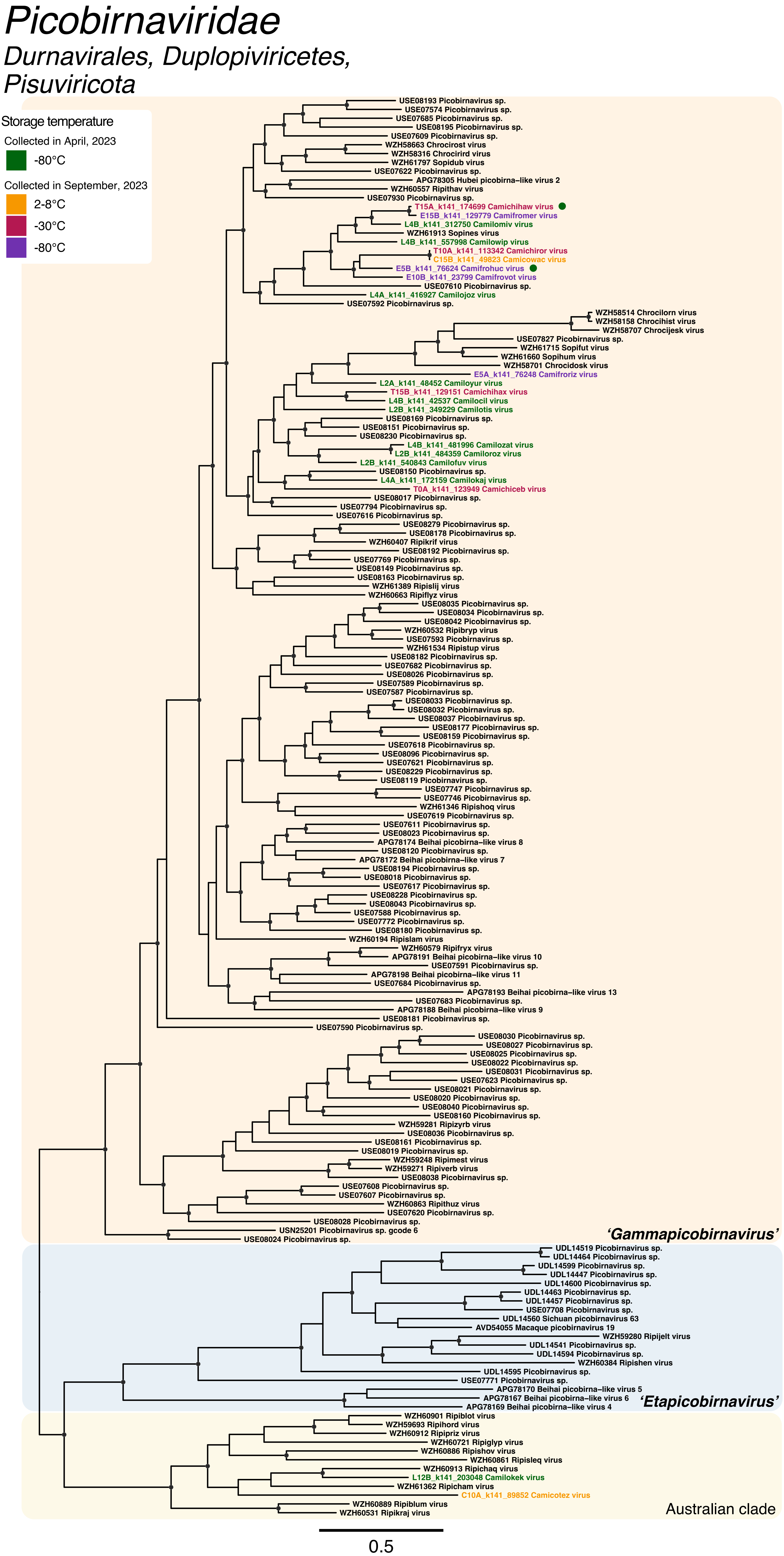
Picobirnaviridae
Durnavirales, Duplopiviricetes, Pisuviricota
Storage temperature
Collected in April, 2023
-80°C
Collected in September, 2023
2-8°C
-30°C
-80°C
USE08193 Picobirnavirus sp.
USE07574 Picobirnavirus sp.
USE07685 Picobirnavirus sp.
USE08195 Picobirnavirus sp.
USE07609 Picobirnavirus sp.
WZH58663 Chrocirost virus
WZH58316 Chrocirird virus
WZH61797 Sopidub virus
USE07622 Picobirnavirus sp.
APG78305 Hubei picobirna−like virus 2
WZH60557 Ripithav virus
USE07930 Picobirnavirus sp.
T15A_k141_174699 Camichihaw virus
E15B_k141_129779 Camifromer virus
L4B_k141_312750 Camilomiv virus
WZH61913 Sopines virus
L4B_k141_557998 Camilowip virus
T10A_k141_113342 Camichiror virus
C15B_k141_49823 Camicowac virus
E5B_k141_76624 Camifrohuc virus
E10B_k141_23799 Camifrovot virus
USE07610 Picobirnavirus sp.
L4A_k141_416927 Camilojoz virus
USE07592 Picobirnavirus sp.
WZH58514 Chrocilorn virus
WZH58158 Chrocihist virus
WZH58707 Chrocijesk virus
USE07827 Picobirnavirus sp.
WZH61715 Sopifut virus
WZH61660 Sopihum virus
WZH58701 Chrocidosk virus
E5A_k141_76248 Camifroriz virus
L2A_k141_48452 Camiloyur virus
T15B_k141_129151 Camichihax virus
L4B_k141_42537 Camilocil virus
L2B_k141_349229 Camilotis virus
USE08169 Picobirnavirus sp.
USE08151 Picobirnavirus sp.
USE08230 Picobirnavirus sp.
L4B_k141_481996 Camilozat virus
L2B_k141_484359 Camiloroz virus
L2B_k141_540843 Camilofuv virus
USE08150 Picobirnavirus sp.
L4A_k141_172159 Camilokaj virus
T0A_k141_123949 Camichiceb virus
USE08017 Picobirnavirus sp.
USE07794 Picobirnavirus sp.
USE07616 Picobirnavirus sp.
USE08279 Picobirnavirus sp.
USE08178 Picobirnavirus sp.
WZH60407 Ripikrif virus
USE08192 Picobirnavirus sp.
USE07769 Picobirnavirus sp.
USE08149 Picobirnavirus sp.
USE08163 Picobirnavirus sp.
WZH61389 Ripislij virus
WZH60663 Ripiflyz virus
USE08035 Picobirnavirus sp.
USE08034 Picobirnavirus sp.
USE08042 Picobirnavirus sp.
WZH60532 Ripibryp virus
USE07593 Picobirnavirus sp.
WZH61534 Ripistup virus
USE08182 Picobirnavirus sp.
USE07682 Picobirnavirus sp.
USE08026 Picobirnavirus sp.
USE07589 Picobirnavirus sp.
USE07587 Picobirnavirus sp.
USE08033 Picobirnavirus sp.
USE08032 Picobirnavirus sp.
USE08037 Picobirnavirus sp.
USE08177 Picobirnavirus sp.
USE08159 Picobirnavirus sp.
USE07618 Picobirnavirus sp.
USE08096 Picobirnavirus sp.
USE07621 Picobirnavirus sp.
USE08229 Picobirnavirus sp.
USE08119 Picobirnavirus sp.
USE07747 Picobirnavirus sp.
USE07746 Picobirnavirus sp.
WZH61346 Ripishoq virus
USE07619 Picobirnavirus sp.
USE07611 Picobirnavirus sp.
USE08023 Picobirnavirus sp.
APG78174 Beihai picobirna−like virus 8
USE08120 Picobirnavirus sp.
APG78172 Beihai picobirna−like virus 7
USE08194 Picobirnavirus sp.
USE08018 Picobirnavirus sp.
USE07617 Picobirnavirus sp.
USE08228 Picobirnavirus sp.
USE08043 Picobirnavirus sp.
USE07588 Picobirnavirus sp.
USE07772 Picobirnavirus sp.
USE08180 Picobirnavirus sp.
WZH60194 Ripislam virus
WZH60579 Ripifryx virus
APG78191 Beihai picobirna−like virus 10
USE07591 Picobirnavirus sp.
APG78198 Beihai picobirna−like virus 11
USE07684 Picobirnavirus sp.
APG78193 Beihai picobirna−like virus 13
USE07683 Picobirnavirus sp.
APG78188 Beihai picobirna−like virus 9
USE08181 Picobirnavirus sp.
USE07590 Picobirnavirus sp.
USE08030 Picobirnavirus sp.
USE08027 Picobirnavirus sp.
USE08025 Picobirnavirus sp.
USE08022 Picobirnavirus sp.
USE08031 Picobirnavirus sp.
USE07623 Picobirnavirus sp.
USE08021 Picobirnavirus sp.
USE08020 Picobirnavirus sp.
USE08040 Picobirnavirus sp.
USE08160 Picobirnavirus sp.
WZH59281 Ripizyrb virus
USE08036 Picobirnavirus sp.
USE08161 Picobirnavirus sp.
USE08019 Picobirnavirus sp.
WZH59248 Ripimest virus
WZH59271 Ripiverb virus
USE08038 Picobirnavirus sp.
USE07608 Picobirnavirus sp.
USE07607 Picobirnavirus sp.
WZH60863 Ripithuz virus
USE07620 Picobirnavirus sp.
USE08028 Picobirnavirus sp.
USN25201 Picobirnavirus sp. gcode 6
USE08024 Picobirnavirus sp.
'Gammapicobirnavirus'
UDL14519 Picobirnavirus sp.
UDL14464 Picobirnavirus sp.
UDL14599 Picobirnavirus sp.
UDL14447 Picobirnavirus sp.
UDL14600 Picobirnavirus sp.
UDL14463 Picobirnavirus sp.
UDL14457 Picobirnavirus sp.
USE07708 Picobirnavirus sp.
UDL14560 Sichuan picobirnavirus 63
AVD54055 Macaque picobirnavirus 19
WZH59280 Ripijelt virus
UDL14541 Picobirnavirus sp.
UDL14594 Picobirnavirus sp.
WZH60384 Ripishen virus
UDL14595 Picobirnavirus sp.
USE07771 Picobirnavirus sp.
APG78170 Beihai picobirna−like virus 5
APG78167 Beihai picobirna−like virus 6
APG78169 Beihai picobirna−like virus 4
'Etapicobirnavirus'
WZH60901 Ripiblot virus
WZH59693 Ripihord virus
WZH60912 Ripipriz virus
WZH60721 Ripiglyp virus
WZH60886 Ripishov virus
WZH60861 Ripisleq virus
WZH60913 Ripichaq virus
L12B_k141_203048 Camilokek virus
WZH61362 Ripicham virus
C10A_k141_89852 Camicotez virus
WZH60889 Ripiblum virus
WZH60531 Ripikraj virus
Australian clade
0.5

**Supplementary Figure 18.** Phylogenetic tree of RdRp amino acid sequences from the family *Picobirnaviridae*. Putative novel viruses identified in this study are coloured by the storage temperature and collection month, such that samples collected in April 2023 (data set 1) and stored at -80°C are coloured in green, while samples collected in September 2023 (data set 2) and stored at 2-8°C are in orange, -30°C in red, and -80°C in purple. Known viruses are coloured in black. Sequences found in both data sets are indicated by a coloured circle next to the provisional virus name, coloured by the storage temperature of library or libraries the identical sequence was found in. Clades comprised exclusively of sequences sourced from Australian environmental samples are indicated by a yellow box. Trees are midpoint rooted with branch lengths scaled according to the number of amino acid substitutions per site. Black circles represent node support ≥80% using 1000 S H-aLRT replicates.

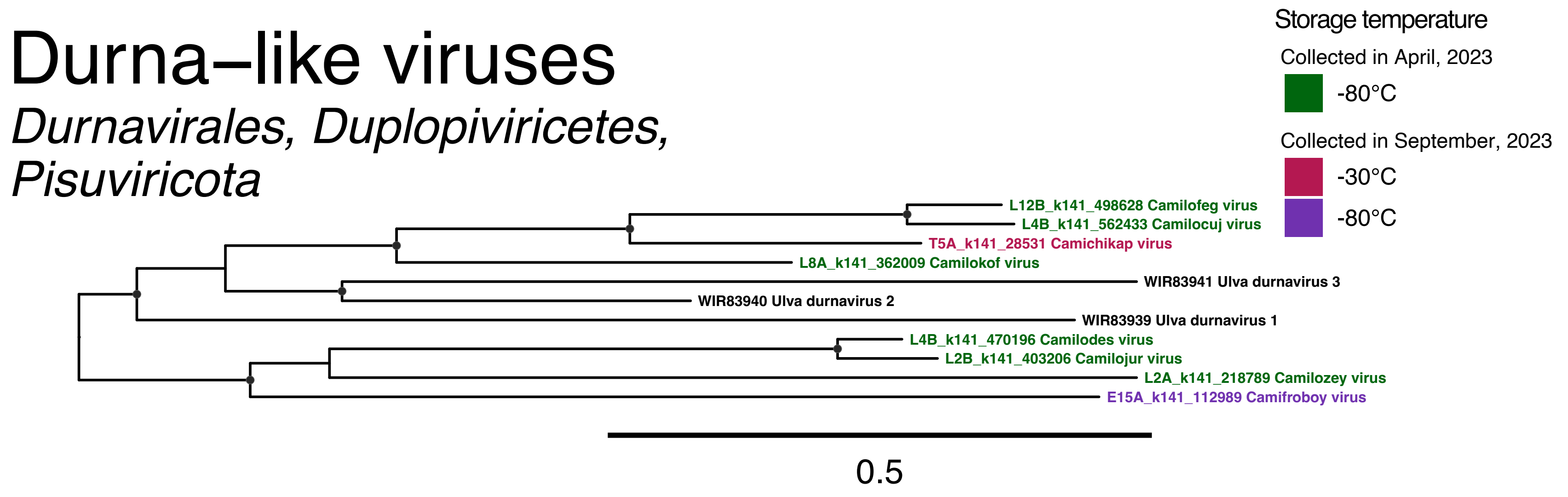
Durna–like viruses
Durnavirales, Duplopiviricetes, Pisuviricota
Storage temperature
Collected in April, 2023
-80°C
Collected in September, 2023
-30°C
-80°C
L12B_k141_498628 Camilofeg virus
L4B_k141_562433 Camilocuj virus
T5A_k141_28531 Camichikap virus
L8A_k141_362009 Camilokof virus
WIR83941 Ulva durnavirus 3
WIR83940 Ulva durnavirus 2
WIR83939 Ulva durnavirus 1
L4B_k141_470196 Camilodes virus
L2B_k141_403206 Camilojur virus
L2A_k141_218789 Camilozey virus
E15A_k141_112989 Camifroboy virus
0.5

**Supplementary Figure 19.** Phylogenetic tree of RdRp amino acid sequences from *Durnavirales*-like sequences. Putative novel viruses identified in this study are coloured by the storage temperature and collection month, such that samples collected in April 2023 (data set 1) and stored at -80°C are coloured in green, while samples collected in September 2023 (data set 2) and stored at -30°C are in red and -80°C in purple. Known viruses are coloured in black. Trees are midpoint rooted with branch lengths scaled according to the number of amino acid substitutions per site. Black circles represent node support ≥80% using 1000 S H-aLRT replicates.

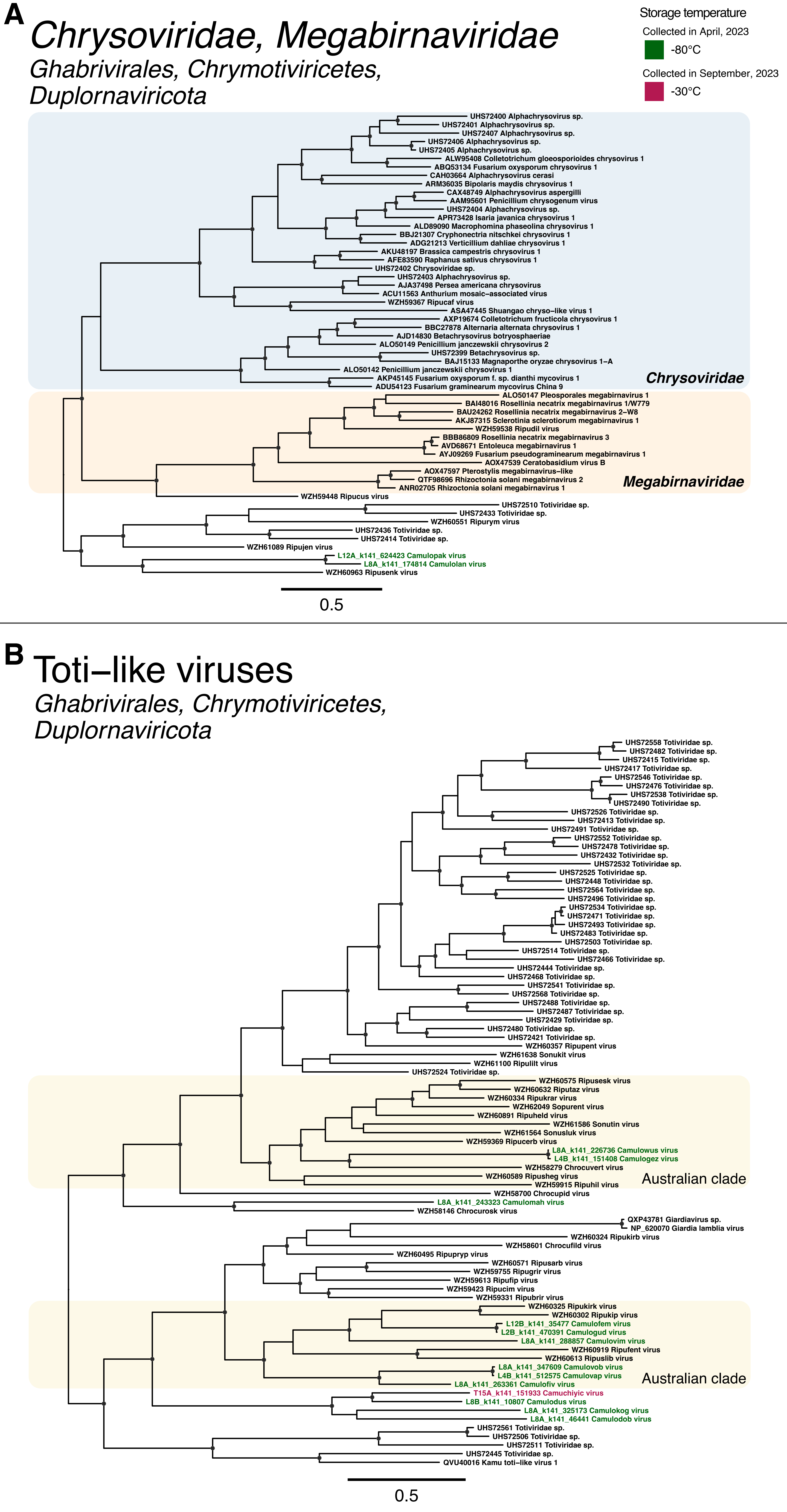
A
Chrysoviridae, Megabirnaviridae
Ghabrivirales, Chrymotiviricetes, Duplornaviricota
Storage temperature
Collected in April, 2023
-80°C
Collected in September, 2023
-30°C
UHS72400 Alphachrysovirus sp.
UHS72401 Alphachrysovirus sp.
UHS72407 Alphachrysovirus sp.
UHS72406 Alphachrysovirus sp.
UHS72405 Alphachrysovirus sp.
ALW95408 Colletotrichum gloeosporioides chrysovirus 1
ABQ53134 Fusarium oxysporum chrysovirus 1
CAH03664 Alphachrysovirus cerasi
ARM36035 Bipolaris maydis chrysovirus 1
CAX48749 Alphachrysovirus aspergilli
AAM95601 Penicillium chrysogenum virus
UHS72404 Alphachrysovirus sp.
APR73428 Isaria javanica chrysovirus 1
ALD89090 Macrophomina phaseolina chrysovirus 1
BBJ21307 Cryphonectria nitschkei chrysovirus 1
ADG21213 Verticillium dahliae chrysovirus 1
AKU48197 Brassica campestris chrysovirus 1
AFE83590 Raphanus sativus chrysovirus 1
UHS72402 Chrysoviridae sp.
UHS72403 Alphachrysovirus sp.
AJA37498 Persea americana chrysovirus
ACU11563 Anthurium mosaic–associated virus
WZH59367 Ripucaf virus
ASA47445 Shuangao chryso–like virus 1
AXP19674 Colletotrichum fructicola chrysovirus 1
BBC27878 Alternaria alternata chrysovirus 1
AJD14830 Betachrysovirus botryosphaeriae
ALO50149 Penicillium janczewskii chrysovirus 2
UHS72399 Betachrysovirus sp.
BAJ15133 Magnaporthe oryzae chrysovirus 1–A
ALO50142 Penicillium janczewskii chrysovirus 1
AKP45145 Fusarium oxysporum f. sp. dianthi mycovirus 1
ADU54123 Fusarium graminearum mycovirus China 9
Chrysoviridae
ALO50147 Pleosporales megabirnavirus 1
BAI48016 Rosellinia necatrix megabirnavirus 1/W779
BAU24262 Rosellinia necatrix megabirnavirus 2–W8
AKJ87315 Sclerotinia sclerotiorum megabirnavirus 1
WZH59538 Ripudil virus
BBB86809 Rosellinia necatrix megabirnavirus 3
AVD68671 Entoleuca megabirnavirus 1
AYJ09269 Fusarium pseudograminearum megabirnavirus 1
AOX47539 Ceratobasidium virus B
AOX47597 Pterostylis megabirnavirus–like
QTF98696 Rhizoctonia solani megabirnavirus 2
ANR02705 Rhizoctonia solani megabirnavirus 1
Megabirnaviridae
WZH59448 Ripucus virus
UHS72510 Totiviridae sp.
UHS72433 Totiviridae sp.
WZH60551 Ripurym virus
UHS72436 Totiviridae sp.
UHS72414 Totiviridae sp.
WZH61089 Ripujen virus
L12A_k141_624423 Camulopak virus
L8A_k141_174814 Camulolan virus
WZH60963 Ripusenk virus
0.5
B
Toti–like viruses
Ghabrivirales, Chrymotiviricetes, Duplornaviricota
UHS72558 Totiviridae sp.
UHS72482 Totiviridae sp.
UHS72415 Totiviridae sp.
UHS72417 Totiviridae sp.
UHS72546 Totiviridae sp.
UHS72476 Totiviridae sp.
UHS72538 Totiviridae sp.
UHS72490 Totiviridae sp.
UHS72526 Totiviridae sp.
UHS72413 Totiviridae sp.
UHS72491 Totiviridae sp.
UHS72552 Totiviridae sp.
UHS72478 Totiviridae sp.
UHS72432 Totiviridae sp.
UHS72532 Totiviridae sp.
UHS72525 Totiviridae sp.
UHS72448 Totiviridae sp.
UHS72564 Totiviridae sp.
UHS72496 Totiviridae sp.
UHS72534 Totiviridae sp.
UHS72471 Totiviridae sp.
UHS72493 Totiviridae sp.
UHS72483 Totiviridae sp.
UHS72503 Totiviridae sp.
UHS72514 Totiviridae sp.
UHS72466 Totiviridae sp.
UHS72444 Totiviridae sp.
UHS72468 Totiviridae sp.
UHS72541 Totiviridae sp.
UHS72568 Totiviridae sp.
UHS72488 Totiviridae sp.
UHS72487 Totiviridae sp.
UHS72429 Totiviridae sp.
UHS72480 Totiviridae sp.
UHS72421 Totiviridae sp.
WZH60357 Ripupent virus
WZH61638 Sonukit virus
WZH61100 Ripulilt virus
UHS72524 Totiviridae sp.
WZH60575 Ripusesk virus
WZH60632 Riputaz virus
WZH60334 Ripukrar virus
WZH62049 Sopurent virus
WZH60891 Ripuheld virus
WZH61586 Sonutin virus
WZH61564 Sonusluk virus
WZH59369 Ripucerb virus
L8A_k141_226736 Camulowus virus
L4B_k141_151408 Camulogez virus
WZH58279 Chrocuvert virus
WZH60589 Ripusheg virus
WZH59915 Ripuhil virus
Australian clade
WZH58700 Chrocupid virus
L8A_k141_243323 Camulomah virus
WZH58146 Chrocurosk virus
QXP43781 Giardiavirus sp.
NP_620070 Giardia lamblia virus
WZH60324 Ripukirb virus
WZH58601 Chrocufild virus
WZH60495 Ripupryp virus
WZH60571 Ripusarb virus
WZH59755 Ripugrir virus
WZH59613 Ripufip virus
WZH59423 Ripucim virus
WZH59331 Ripubrir virus
WZH60325 Ripukirk virus
WZH60302 Ripukip virus
L12B_k141_35477 Camulofem virus
L2B_k141_470391 Camulogud virus
L8A_k141_288857 Camulovim virus
WZH60919 Ripufent virus
WZH60613 Ripuslib virus
L8A_k141_347609 Camulovob virus
L4B_k141_512575 Camulovap virus
L8A_k141_263361 Camulofiv virus
Australian clade
T15A_k141_151933 Camuchiyic virus
L8B_k141_10807 Camulodus virus
L8A_k141_325173 Camulokog virus
L8A_k141_46441 Camulodob virus
UHS72561 Totiviridae sp.
UHS72506 Totiviridae sp.
UHS72511 Totiviridae sp.
UHS72445 Totiviridae sp.
QVU40016 Kamu toti–like virus 1
0.5

**Supplementary Figure 20.** Phylogenetic tree of RdRp amino acid sequences from (A) the families *Chrysoviridae* and *Megabirnaviridae*, and (B) divergent toti-like sequences. Putative novel viruses identified in this study are coloured by the storage temperature and collection month, such that samples collected in April 2023 (data set 1) and stored at -80°C are coloured in green, while samples collected in September 2023 (data set 2) and stored at -30°C are in red. Known viruses are coloured in black. Clades comprised exclusively of sequences sourced from Australian environmental samples are indicated by a yellow box. Trees are midpoint rooted with branch lengths scaled according to the number of amino acid substitutions per site. Black circles represent node support ≥80% using 1000 S H-aLRT replicates.

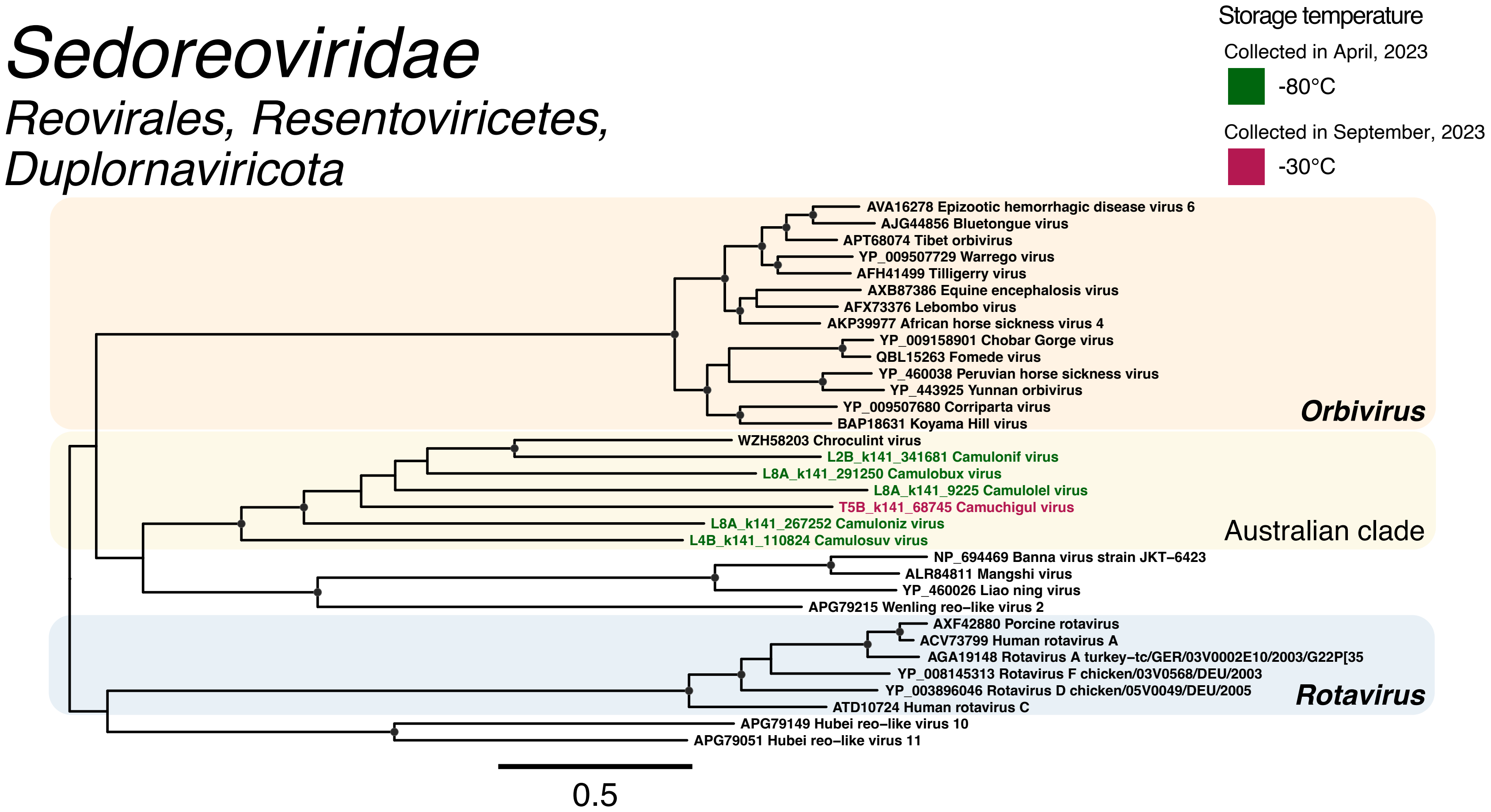

Sedoreoviridae
Reovirales, Resentoviricetes, Duplornaviricota
Storage temperature
Collected in April, 2023
-80°C
Collected in September, 2023
-30°C
AVA16278 Epizootic hemorrhagic disease virus 6
AJG44856 Bluetongue virus
APT68074 Tibet orbivirus
YP_009507729 Warrego virus
AFH41499 Tilligerry virus
AXB87386 Equine encephalosis virus
AFX73376 Lebombo virus
AKP39977 African horse sickness virus 4
YP_009158901 Chobar Gorge virus
QBL15263 Fomede virus
YP_460038 Peruvian horse sickness virus
YP_443925 Yunnan orbivirus
YP_009507680 Corriparta virus
BAP18631 Koyama Hill virus
Orbivirus
WZH58203 Chroculint virus
L2B_k141_341681 Camulonif virus
L8A_k141_291250 Camulobux virus
L8A_k141_9225 Camulolel virus
T5B_k141_68745 Camuchigul virus
L8A_k141_267252 Camuloniz virus
L4B_k141_110824 Camulosuv virus
Australian clade
NP_694469 Banna virus strain JKT–6423
ALR84811 Mangshi virus
YP_460026 Liao ning virus
APG79215 Wenling reo–like virus 2
AXF42880 Porcine rotavirus
ACV73799 Human rotavirus A
AGA19148 Rotavirus A turkey–tc/GER/03V0002E10/2003/G22P[35
YP_008145313 Rotavirus F chicken/03V0568/DEU/2003
YP_003896046 Rotavirus D chicken/05V0049/DEU/2005
ATD10724 Human rotavirus C
Rotavirus
APG79149 Hubei reo–like virus 10
APG79051 Hubei reo–like virus 11
0.5

**Supplementary Figure 21.** Phylogenetic tree of RdRp amino acid sequences from the family *Sedoreoviridae*. Putative novel viruses identified in this study are coloured by the storage temperature and collection month, such that samples collected in April 2023 (data set 1) and stored at -80°C are coloured in green, while samples collected in September 2023 (data set 2) and stored at -30°C are in red. Known viruses are coloured in black. Trees are midpoint rooted with branch lengths scaled according to the number of amino acid substitutions per site. Black circles represent node support ≥80% using 1000 S H-aLRT replicates.

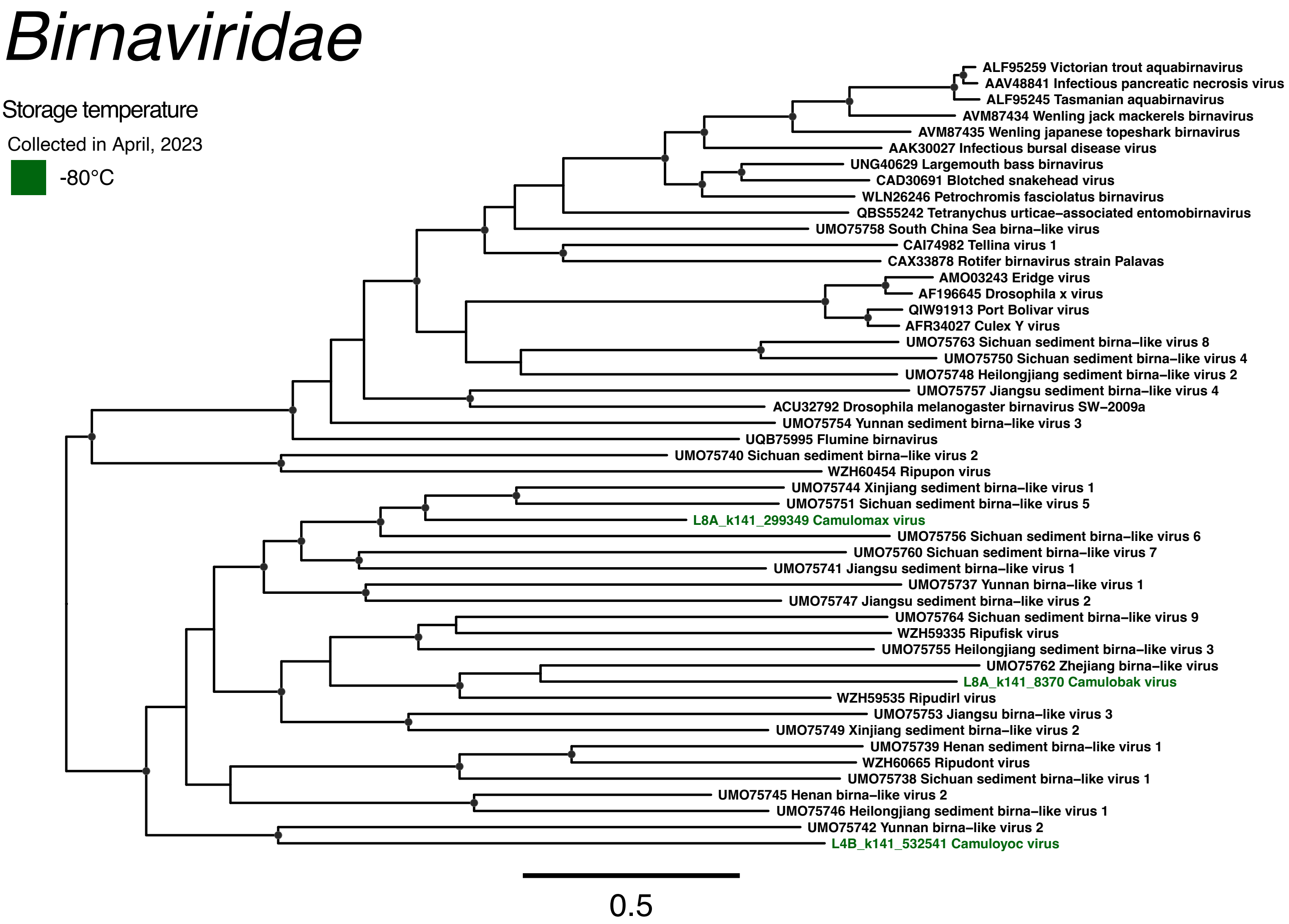
Birnaviridae
Storage temperature
Collected in April, 2023
-80°C
ALF95259 Victorian trout aquabirnavirus
AAV48841 Infectious pancreatic necrosis virus
ALF95245 Tasmanian aquabirnavirus
AVM87434 Wenling jack mackerels birnavirus
AVM87435 Wenling japanese topeshark birnavirus
AAK30027 Infectious bursal disease virus
UNG40629 Largemouth bass birnavirus
CAD30691 Blotched snakehead virus
WLN26246 Petrochromis fasciolatus birnavirus
QBS55242 Tetranychus urticae–associated entomobirnavirus
UMO75758 South China Sea birna–like virus
CAI74982 Tellina virus 1
CAX33878 Rotifer birnavirus strain Palavas
AMO03243 Eridge virus
AF196645 Drosophila x virus
QIW91913 Port Bolivar virus
AFR34027 Culex Y virus
UMO75763 Sichuan sediment birna–like virus 8
UMO75750 Sichuan sediment birna–like virus 4
UMO75748 Heilongjiang sediment birna–like virus 2
UMO75757 Jiangsu sediment birna–like virus 4
ACU32792 Drosophila melanogaster birnavirus SW–2009a
UMO75754 Yunnan sediment birna–like virus 3
UQB75995 Flumine birnavirus
UMO75740 Sichuan sediment birna–like virus 2
WZH60454 Ripupon virus
UMO75744 Xinjiang sediment birna–like virus 1
UMO75751 Sichuan sediment birna–like virus 5
L8A_k141_299349 Camulomax virus
UMO75756 Sichuan sediment birna–like virus 6
UMO75760 Sichuan sediment birna–like virus 7
UMO75741 Jiangsu sediment birna–like virus 1
UMO75737 Yunnan birna–like virus 1
UMO75747 Jiangsu sediment birna–like virus 2
UMO75764 Sichuan sediment birna–like virus 9
WZH59335 Ripufisk virus
UMO75755 Heilongjiang sediment birna–like virus 3
UMO75762 Zhejiang birna–like virus
L8A_k141_8370 Camulobak virus
WZH59535 Ripudirl virus
UMO75753 Jiangsu birna–like virus 3
UMO75749 Xinjiang sediment birna–like virus 2
UMO75739 Henan sediment birna–like virus 1
WZH60665 Ripudont virus
UMO75738 Sichuan sediment birna–like virus 1
UMO75745 Henan birna–like virus 2
UMO75746 Heilongjiang sediment birna–like virus 1
UMO75742 Yunnan birna–like virus 2
L4B_k141_532541 Camuloyoc virus
0.5

**Supplementary Figure 22.** Phylogenetic tree of RdRp amino acid sequences from the family *Birnaviridae*. Putative novel viruses identified in this study are coloured by the storage temperature and collection month, such that samples collected in April 2023 (data set 1) and stored at -80°C are coloured in green. Known viruses are coloured in black. Clades comprised exclusively of sequences sourced from Australian environmental samples are indicated by a yellow box. Trees are midpoint rooted with branch lengths scaled according to the number of amino acid substitutions per site. Black circles represent node support ≥80% using 1000 S H-aLRT replicates.
